# Reproductive isolation via polygenic local adaptation in sub-divided populations: effect of linkage disequilibria and drift

**DOI:** 10.1101/2021.11.05.467433

**Authors:** Himani Sachdeva

**Affiliations:** Department of Mathematics, University of Vienna, Vienna, Austria

## Abstract

This paper considers how polygenic local adaptation and reproductive isolation between hybridizing populations is influenced by linkage disequilibria (LD) between loci, in scenarios where both gene flow and genetic drift counteract selection. It shows that the combined effects of multi-locus LD and genetic drift on allele frequencies at selected loci and on heterozygosity at neutral loci are predicted accurately by incorporating (deterministic) effective migration rates into the diffusion approximation (for selected loci) and into the structured coalescent (for neutral loci). Theoretical approximations are tested against individual-based simulations and used to investigate conditions for the maintenance of local adaptation on an island subject to one-way migration from a differently adapted mainland, and in an infinite-island population with two habitats under divergent selection. The analysis clarifies the conditions under which LD between sets of locally deleterious alleles allows these to be collectively eliminated despite drift, causing sharper and (under certain conditions) shifted migration thresholds for loss of adaptation. Local adaptation also has counter-intuitive effects on neutral (relative) divergence: *F*_*ST*_ is highest for a pair of subpopulations belonging to the same (rare) habitat, despite the lack of reproductive isolation between them.

**Author summary:** Environmental adaptation often involves spatially heterogeneous selection at many genetic loci. Thus, the evolutionary consequences of hybridisation between populations adapted to different environments depend on the *coupled* dynamics of multiple loci under selection, migration and genetic drift, making them challenging to predict. Here, I introduce theoretical approximations that accurately capture the effect of such coupling on allele frequencies at individual loci, while also accounting for the stochastic effects of genetic drift. I then use these approximations to study hybridisation in a metapopulation consisting of many interconnected subpopulations, where each subpopulation belongs to one of two habitats under divergent selection. The analysis clarifies how subpopulations belonging to a rare habitat can maintain local adaptation despite high levels of migration if net selection against multi-locus genotypes is stronger than a threshold which depends on the relative abundances of the two habitats. Further, local adaptation in a metapopulation can significantly elevate *F*_*ST*_ between subpopulations belonging to the same habitat, even though these are not reproductively isolated. These findings highlight the importance of carefully considering the genetic architecture and spatial context of divergence when interpreting patterns of genomic differentiation between speciating populations.

## Introduction

Environmental adaptation in natural populations typically involves selection that varies over space and time, and acts on many genetic loci [1–4]. The ability of populations to adapt to their local environment thus depends on the ease with which locally favoured alleles can establish and be maintained across multiple loci, despite maladaptive gene flow from other regions. A key question is: to what extent does selection act on *combinations* of selected alleles, as opposed to individual alleles [5, 6]? More generally, when do linkage disequilibria (LD), i.e., statistical associations between sets of locally adaptive alleles— that may be either tightly clustered or widely distributed across the genome— protect such alleles from swamping, especially in marginal habitats, which are prone to swamping [7]?

The buildup and/or maintenance of LD between sets of divergently selected alleles may be viewed as an example of the broader process of ‘coupling’ between barrier loci, which reduces genetic exchange between populations [8, 9]— a precursor to reproductive isolation (RI) and speciation. However, many questions remain as to the role of such coupling during the initial vs. late stages of divergence, its role in allopatric vs. parapatric divergence, and in case of the latter, whether divergence (with gene flow) involves adaptation from standing genetic variation or new mutations [10].

Barton (1983) [5] considered the consequences of secondary contact between populations subject to divergent directional selection at multiple equal-effect loci, and showed that in large populations, where the effects of drift can be neglected, the strength of LD between a set of introgressing deleterious alleles depends on the selection density, i.e., the ratio of the total selection strength (against the entire set of alleles) to the total recombination rate (over the map length spanned by the alleles). If total selection is stronger than total recombination, and immigration sufficiently weak that deleterious alleles segregate at low frequencies, *sets* of such alleles are eliminated by selection much faster than they are broken apart by recombination. The frequency of any allele at migration-selection equilibrium is then influenced more by indirect selection due to LD with other deleterious alleles than direct selection due to its own deleterious effect. Conversely, when recombination is much faster than selection, allele frequencies evolve more or less independently across loci, primarily under direct selection.

Selection against introgressing deleterious genotypes also impedes neutral gene flow if neutral alleles are in strong LD with alleles that are divergently selected across populations. This reduction in neutral exchange can be quantified in terms of *effective* migration rates (in case of discrete populations connected via migration; see e.g., [11]) or the strength of a ‘barrier’ to gene flow (in case of spatially continuous populations subject to heterogeneous selection; e.g., [12]). Barton and Bengtsson (1986) [13] calculated effective migration rates and barrier strengths in large populations (i.e., neglecting drift) for a variety of multilocus configurations and spatial geometries. They showed that when barrier loci are linked across a linear genome, then the effective migration rate of a neutral allele at an arbitrary genomic location is strongly reduced only when selection density, i.e, net selection against introgressing deleterious alleles per unit map length, is high. With unlinked barrier loci, effective migration rates at neutral markers are reduced relative to the actual migration rate by a factor that depends (to a good approximation) only on the relative fitness of immigrants [11].

A more complex picture emerges for divergence with ongoing gene flow— a locally beneficial mutation that arises near a pre-existing barrier locus enjoys increased chances of establishment (so-called ‘divergence hitchhiking’; see e.g., [14]). However, this effect is typically restricted to a small map region (with a correspondingly small mutation target) around the barrier locus, and thus, does not markedly influence the rate of buildup of divergence between populations. If the genomic density of barrier loci is low (as in early stages of divergence), any newly arisen mutation will find itself in the vicinity of at most one such locus, allowing us to investigate the effects of divergence hitchhiking on the establishment of new mutations using relatively simple two-locus models [15–19].

As divergence builds up, any new mutation comes to be influenced by more barrier loci- until a critical divergence threshold is reached, beyond which there is a sharp decline in effective migration rates and a corresponding increase in establishment probabilities of locally adaptive variants across the *entire* genome [20, 21]. This has been hypothesised to cause a ‘tipping point’ in the course of speciation, with the buildup of RI rapidly accelerating beyond this point [22]. However, an important limitation of these studies is that they only consider divergence via the sequential establishment of new mutations (as opposed to response from standing genetic variation). Moreover, they rely largely on simulations, making it difficult to arrive at a complete picture of multilocus interactions during adaptive divergence.

A further complication arises when populations are small and drift comparable to selection per locus. Not only can drift, in conjunction with maladaptive gene flow, then impair the efficacy of selection at individual loci [23], it might also generate negative LD between sets of deleterious alleles via Hill-Robertson interference [24]: this may counteract positive LD due to migration, thus also compromising the efficacy of LD-driven or collective elimination of groups of alleles.

Understanding the combined effects of LD and genetic drift on local adaptation is important as extended populations are often patchworks of smaller, interconnected subpopulations. If density regulation occurs primarily within subpopulations, then the rate of drift is governed by local sizes rather than the size of the population as a whole. Additionally, if locally adaptive traits are polygenic, then individual loci contributing to trait variation may have rather weak selective effects, such that typical values of *Ns* are small, resulting in local adaptation via many small (and possibly transient) allele frequency shifts [25, 26].

This paper considers some of these issues by analysing polygenic local adaptation in a metapopulation comprised of many small subpopulations occupying different habitats, assuming that fitness is influenced by many loci with habitat-dependent selective effects. It explores conditions for local adaptation in two scenarios— first, for an island subject to maladaptive gene flow from a large and perfectly adapted mainland, and second, in an infinite-island population with two habitats subject to divergent selection. The focus is on understanding when LD between locally adaptive alleles allows adaptation to be maintained in a *rare* habitat (which encompasses a small fraction of all islands) despite migration, and to what extent the effects of LD may be washed out by drift. A key question is: How does the genetic architecture of local adaptation (i.e., the number and effect sizes of locally adaptive variants) influence evolutionary outcomes in a scenario where adaptation involves response from high levels of standing genetic variation? I further explore how neutral diversity in either habitat is influenced by the extent of adaptive divergence, and how this translates into expectations for various *F*_*ST*_ measures.

The paper also illustrates how the gross effects of multi-locus LD and genetic drift at any individual selected locus are accurately predicted by incorporating appropriately defined effective migration rates (for selected alleles) into the single-locus diffusion approximation (see e.g., [27]). Analogously, the effects of LD on neutral diversity within any subpopulation are captured by incorporating effective migration rates (for neutral alleles) into the structured coalescent. While the basic approach of splicing effective migration rates into the single-locus diffusion approximation has been employed in earlier work on two-locus models [28–32], as we see below, this approach turns out to be especially powerful when many loci (spread across the entire genome) are involved in divergence: the effective migration rate at any locus is then roughly independent of its own effect, depending instead on the relative fitness of migrant individuals, which can often be estimated in the field, e.g., in reciprocal transplant experiments [33], or from pedigrees [34].

More generally, a mathematical understanding of multi-locus evolution under selection and drift remains elusive, despite the centrality of such an understanding to fundamental evolutionary questions regarding the limits to natural selection, the evolution of sex and recombination, and the maintenance of genetic variation. Thus, heuristic approximations (of the kind developed here) can play an important role in our understanding of stochastic effects during polygenic adaptation [33, 35–37].

## Models and Methods

### Mainland-island model

Consider an island with *N* haploid individuals, subject to ongoing migration from a large mainland. Island and mainland populations are under divergent selection at *L* unlinked, biallelic loci. Selection is multiplicative across loci, with different alleles favoured on the mainland and island at each locus, independent of the state of other loci. For simplicity, we will also take effect sizes to be equal across loci. Then, the fitness of any individual on the island depends only on *y*, the number of locally deleterious alleles it carries, and *s*, the selective effect per deleterious allele, and is given by *W* (*y*)=*e*^*−sy*^.

We assume that the mainland population is fixed at all selected loci for the allele that is deleterious on the island, so that immigrants have fitness proportional to *e*^*−sL*^. Polymorphism can still be maintained on the island despite drift and continual gene flow from the mainland, provided mutation rates are above some threshold value.

The lifecycle on the island is as follows: in each generation, a Poisson-distributed number of individuals (on average *Nm*) are replaced by migrants from the mainland, where *m* is the migration rate. Individuals then undergo mutation, with rate of mutation *µ* between alternative alleles per locus per individual. Following mutation, the next generation is formed by sampling 2*N* parents (with replacement) with probabilities proportional to their relative fitness. Finally, the 2*N* parents are paired randomly and recombinant haploid offspring created via free recombination between pairs.

### Infinite-island model

Consider a population with *D* islands, where each island has *N* haploid individuals. In theoretical analyses, we will assume *D* → ∞, i.e., consider the infinite-island model [27]. The fitness of an individual depends on the local environment or *habitat* on the island on which it resides. For simplicity, we will assume only two habitats, with a fraction *ρ* of islands supporting the first habitat, and the remaining fraction 1 − *ρ* the second. Assuming *ρ<*1*/*2, the first habitat is always ‘rare’ (i.e., it characterizes a minority of islands in the population) and the second ‘common’.

As before, fitness is influenced by *L* unlinked, biallelic, equal-effect loci, with alternative alleles favoured in the two habitats at each locus. The relative fitness of an individual carrying *i locally* deleterious alleles in a deme belonging to habitat *k* is 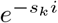, where *s*_*k*_ is the selective effect per locally deleterious allele in habitat *k*. We will use the subscripts *r* and *c* to denote the rare and common habitats. For simplicity, we will only consider symmetric selection (with *s*_*r*_=*s*_*c*_=*s*), but the approximations described below apply more generally.

With an infinite number of demes, the mutation rate can be set to zero, since mutation is not essential for the maintenance of polymorphism as long as *ρ* is not vanishingly small, and because we are primarily interested in the effect of gene flow on local adaptation from standing variation. The extension to non-zero mutation rates is straightforward.

In each generation and in each deme, a Poisson-distributed number of individuals (with mean *Nm*) are replaced by migrants from a common pool, which is formed by drawing individuals uniformly from across all demes. Following migration, the next generation is formed by randomly sampling 2*N* parents (within each deme) with probabilities proportional to local relative fitness. *N* haploid offspring are then created in each deme by free recombination between parental pairs. For the infinite-island model, we will also follow neutral markers that are unlinked to any selected locus and to each other— this allows us to investigate how the extent of adaptive divergence between habitats influences neutral gene flow and genome-wide RI.

Since the main goal is to clarify how LD and drift jointly influence local adaptation in a structured population, other kinds of complexity are neglected. Organisms are assumed to be haploid (thus neglecting the effects of dominance). Loci are assumed to be unlinked and selective effects taken to be the same for all loci. More crucially, the model assumes an extreme form of divergent selection, wherein any selected allele has opposite effects in the two habitats (regardless of alleles at other loci). Finally, there is no explicit space and no isolation-by-distance.

In the following, I briefly outline theoretical approximations that predict allele frequency divergence in different limiting cases: I first discuss the single-locus diffusion approximation for allele frequencies in a subdivided population [27]– this accounts for the effects of genetic drift but neglects LD. I then discuss deterministic analyses that account for LD but neglect drift [5, 13]; it is useful to represent the effects of LD in such analyses by an effective migration rate for the selected or neutral allele [11, 28]. Finally, I describe how effective migration rates can be incorporated into the diffusion approximation, leading to novel approximations that accurately predict allele frequency divergence in parameter regimes where both multi-locus LD and drift play a role.

### Diffusion approximation (assuming LE)

If net selection against maladapted genotypes is weak relative to recombination, then LD between selected variants can be neglected and loci assumed to evolve independently, i.e., under linkage equilibrium (LE). For definiteness, we will use *p* to denote the frequency of the allele that is locally disadvantageous in the rare habitat (or in case of mainland-island migration, on the island) and thus advantageous in the common habitat (or on the mainland, where we assume *p*=1). If 1*/N, s, m, µ* « 1, then the probability distribution *ψ*[*p*] of the allele frequency *p* at any locus under mutation-selection-migration-drift equilibrium is predicted by the diffusion approximation, and depends only on the scaled parameters *Ns, Nµ* and *Nm* [27].

The equilibrium frequency distribution on an island subject to one-way migration from the mainland is given by (see e.g., [27] for details):

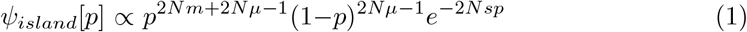

Integrating over the normalised distribution gives the expected allele frequency 𝔼 [*p*].

For the infinite-island model, one can express the allele frequency distribution 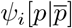 at any locus in a deme within habitat *i*, as a function of 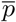, the allele frequency at that locus in the migrant pool [27]:

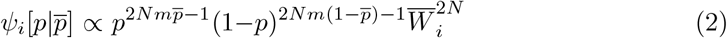

where 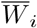 is the mean population fitness for a deme in habitat *i*. The subscript *i* can take on values *r* and *c*, corresponding to the rare and common habitats respectively. We have: 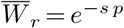 and 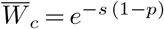, since alternative alleles are favoured in the two habitats. One can now calculate the expected allele frequency 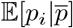 in either habitat as a function of 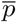 by integrating over the frequency distribution above. At equilibrium, 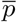 must be equal to the expected allele frequency across the entire population. This allows us to obtain 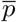 by numerically solving 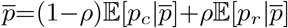 (see also [38]).

### Effective migration rates (neglecting drift)

The assumption of LE is valid only if recombination is faster than all other evolutionary processes. In particular, this requires *Ls* « 1*/*2, i.e., the net selective disadvantage of maladapted immigrant genotypes must be much weaker than recombination. Conversely, with strong selection against immigrants i.e., *Ls* ≳ 1*/*2, sets of incoming alleles are eliminated together before recombination can split them, causing allele frequencies across different loci to evolve in a coupled manner. Thus, in this regime, we must explicitly consider multilocus dynamics in order to account for the effects of LD on allele frequencies [5].

Consider a large population that receives migrants, carrying *L* unlinked deleterious alleles, at a steady rate *m* per generation. Let { *P*_*y*_ } denote the frequencies of genotypes with *y*=1, 2, … *L* deleterious alleles in the population at equilibrium. If selection per locus and migration are much stronger than drift, i.e., 1*/N* « *m* ∼ *s* « *Ls* ∼ 1, then genotype frequencies evolve essentially deterministically. Further, if deleterious genotypes are rare at equilibrium (e.g., as expected for large *Ls*), then mating between individuals both carrying deleterious genotypes can be neglected. Under these conditions, {*P*_*y*_} satisfy the following coupled *linear* equations:

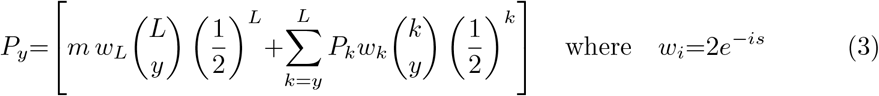

Here, *w*_*k*_=2*e*^*−ks*^ is the average number of offspring of an individual carrying *k* deleterious alleles, and 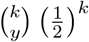 the probability that the individual transmits exactly *y* of these deleterious alleles to an offspring. Note that for *s*=0, individuals have an average of 2 offspring and transmit half of their genome to any offspring on average.

Eq (3) can be solved to obtain the average deterministic deleterious allele frequency 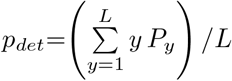 (Section 1, S1 Text). One can then use this to define the effective migration rate *m*_*e*_[*s, L*] for a selected allele as that rate of migration which would cause the allele frequency at a *single* locus under migration-selection balance (with selective disadvantage *s* for the deleterious allele) to be the same as *p*_*det*_, the average deleterious frequency that emerges in the multi-locus model, where immigrant genotypes carrying *L* such deleterious alleles are introduced at rate *m* per generation. Then, we have (see Section 1, S1 Text):

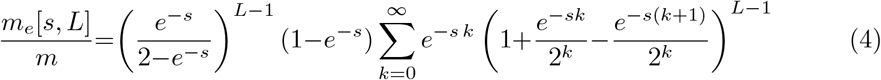

Following [13], we can also calculate an effective migration rate 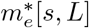 for neutral alleles. By definition, 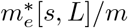 is the probability that the neutral allele escapes via one or more recombination events from the immigrant genetic background (which has an excess of *L* deleterious alleles of effect *s*) onto a resident background, before it is lost from the population. For an unlinked neutral allele, this is (Section 1, S1 Text):

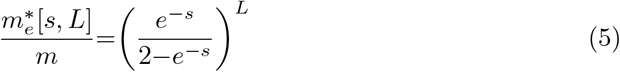

In the limit *s* → 0, *L* →∞, with *Ls* constant, i.e., assuming that a given total selective disadvantage *Ls* is due to larger and larger numbers of loci of weaker effects, the two (scaled) effective migration rates, *m*_*e*_[*s, L*]*/m* and 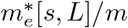, can be approximated as (see Section 1, S1 Text):

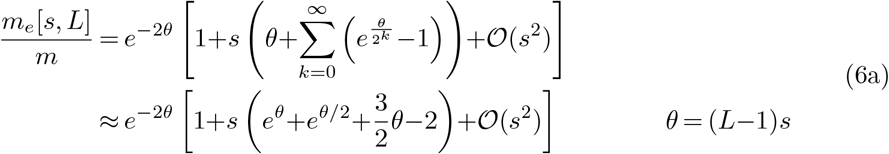

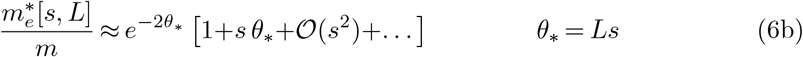

Note that in this limit, we also have *θ* → *θ*_*∗*_. However, I still distinguish between the two to highlight the conceptual distinction that the barrier effect at any selected locus is due to the other *L* − 1 selected loci, while the barrier effect at a neutral locus is due to *L* selected loci. Figure 1 in S1 Text illustrates how the ratios *m*_*e*_[*s, L*]*/m* and 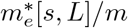 converge towards the large-*L*/small-*s* predictions of Eq (6) for various values of *Ls*.

**Fig 1.**
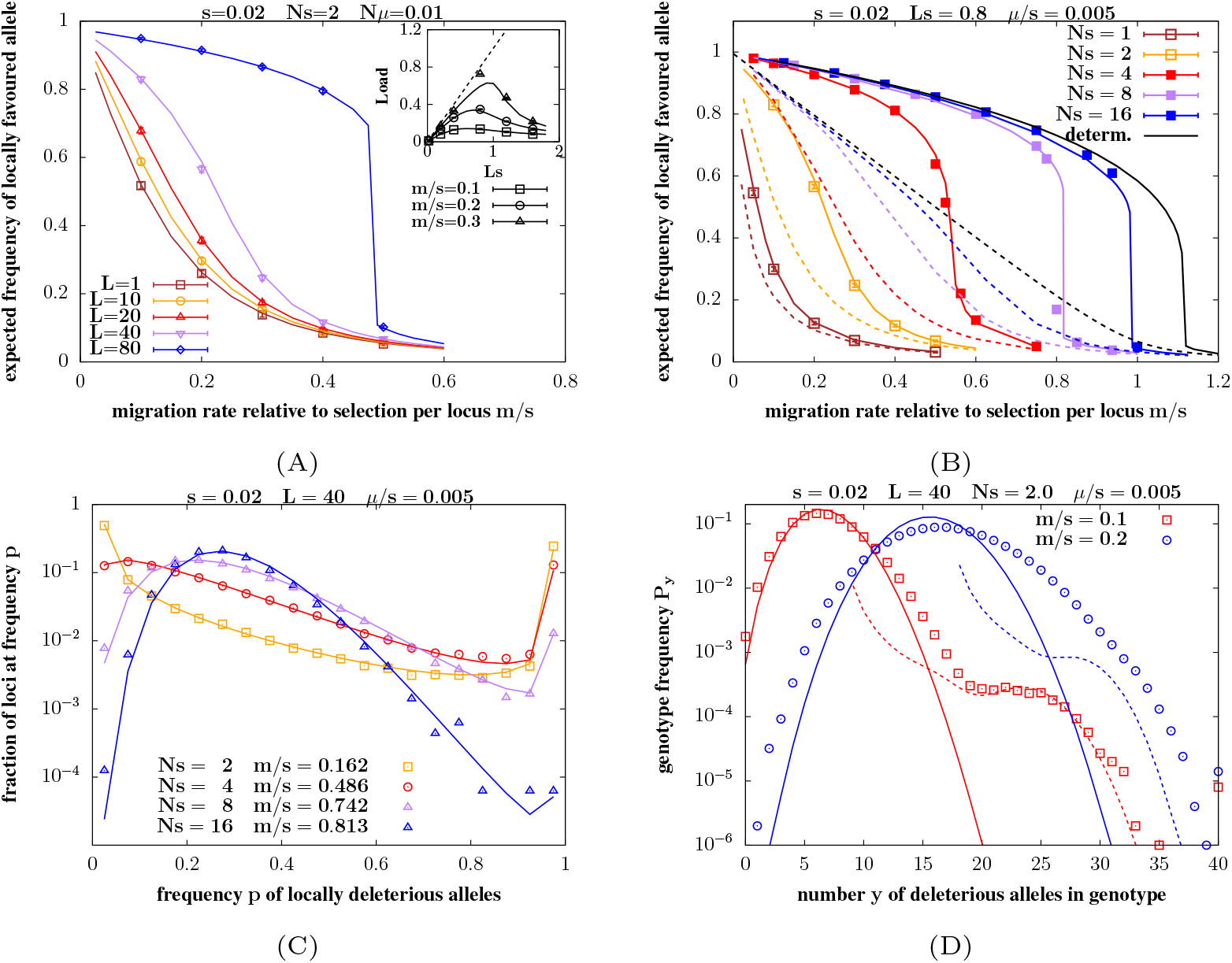
Local adaptation under mainland-island migration. A. Expected frequency 1 − E[*p*] of the locally favoured allele on the island vs. *m/s*, the migration rate relative to selection per locus, for various *L* (different colors) for *s*=0.02, *Ns*=2, and *µ/s*=0.005. Inset: Expected load vs. *Ls* (which is varied by changing *L*) for various values of *m/s*. The maximum possible load *Ls* (dashed line) is also shown for reference. B. Expected frequency of the locally favoured allele vs. *m/s* for various *Ns* (different colors) for *s*=0.02, *L*=40 and *µ/s*=0.005. Symbols depict results of individual-based simulations in both Figs 1A and 1B (obtained by averaging over 100 − 200 simulation replicates for each point). Colored solid lines show theoretical predictions that account for both LD and drift (obtained from Eq (7) together with Eq (4)); colored dashed lines in 1B show LE/single-locus predictions that only account for drift (and are obtained from Eq (1)). Fig 1B also shows deterministic predictions that account for LD (solid black line) as well as the LE/single-locus deterministic prediction (dashed black line). See main text for how these are calculated. Note that there are no simulation results for the deterministic case (as individual-based simulations are always affected by drift). C. Distribution of allele frequencies shown by plotting the fraction of loci with frequency of locally deleterious allele between *p* and *p*+Δ*p*, vs. *p* (for Δ*p*=0.05). The different colors show distributions for *N* =100, 200, 400, 800 (which correspond to *Ns*=2, 4, 8, 16, for *s*=0.02), with *m/s* chosen in each case such that the expected frequency of the locally deleterious allele is 0.3. Theoretical allele frequency distributions (lines) match well with those from individual-based simulations (symbols), with some (moderate) deviation in larger populations. Theoretical predictions are obtained using Eq (1) with *m* replaced by *m*_*e*_, which depends on E[*p*], which is determined numerically, as above. D. Genotype frequencies *P*_*y*_, which represent the probability that a randomly chosen genotype in the population carries *y* deleterious alleles, vs. *y*, for two different values of *m/s*, for *Ns*=2 and *s*=0.02. Symbols depict results of individual-based simulations; solid lines show deterministic predictions and dashed lines predictions under LE (see text for more details about the two kinds of predictions). Other parameters for C. and D. are: *L*=40 and *µ/s*=0.005.

We can see from Eq (6) (see also Figure 1 in S1 Text) that *m*_*e*_*/m* is lowest or the barrier effect *strongest* when the same total selective disadvantage *Ls* is due to a very large number of loci of very weak effect. In this limit, the effective migration rate of any allele (neutral or selected) is reduced relative to *m* by approximately *e*^*−*2*θ*^, where *θ* is the net selective disadvantage (in the recipient population) of the genetic background of the immigrating allele. One can also arrive at this result in a less rigorous but more general way from the fact that 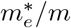 for a neutral unlinked allele (in the limit of weak migration) is equal to the average *reproductive value* (RV) of migrants [39]. Here, RV refers to the migrant’s long-term genetic contribution to the recipient population [40]. An approximate expression for the RV of the migrant in the highly polygenic limit can be derived as follows (see also [11]).

Let *K* denote the average number of locally deleterious alleles per genome in the resident population. Migrants carry an excess of *L* − *K* deleterious alleles with respect to the average resident, and their relative fitness is *e*^*−*(*L−K*)*s*^ (neglecting fitness variance within the resident population). Since the immediate progeny of the migrant (i.e., *F*_1_ individuals) will carry 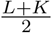 deleterious alleles on average, i.e., an excess of 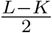 deleterious alleles, their relative fitness is ≈ *e*^*−s* (*L−K*)*/*2^. For *m* « 1 (i.e., when mating between individuals with recent immigrant ancestry can be neglected), most second-generation descendants of the immigrant individual are first-generation backcrosses; thus, they carry an excess of 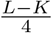 deleterious alleles on average and have relative fitness ≈ *e*^*−s* (*L−K*)*/*4^. Similarly, third-generation descendants would have fitness ≈ *e*^*−s* (*L−K*)*/*8^, and so on. Thus, the average RV of a migrant, which is the product of its own relative fitness with that of all its descendants, is approximately *e*^*−s*(*L−K*)^ *e*^*−s*(*L−K*)*/*2^ *e*^*−s*(*L−K*)*/*4^ *e*^*−s*(*L−K*)*/*8^ … = *e*^*−*2*s*(*L−K*)^. Conceptually similar arguments (based on tracking the long-term genetic contribution of individuals in a population) have also been used to derive how fitness variance at unlinked loci affects effective population size [41].

Note that we recover *m*_*e*_*/m* ≈ *e*^*−*2*Ls*^ (as in Eq (6) above) for very rare migration, i.e., when most genotypes in the resident population have no deleterious alleles (*K* ≈ 0). This derivation is only approximate as it neglects the segregation variance among descendants of the migrant individual as well as fitness variance in the resident population. The derivations based on Eq (3) (see also eq. 6 in S1 Text) account for the former by summing over the frequencies of all possible offspring genotypes. However, this only leads to corrections that are 𝒪 (*s*) and are thus small for weak per-locus selection.

### Introducing effective migration rates into the diffusion approximation

If individual subpopulations are small, drift may be comparable to selection per deleterious allele, but much weaker than selection against immigrant genotypes (that carry many such alleles). This corresponds to a parameter regime with 1*/N* ∼ *m* ∼ *s* « *Ls* ∼ 1. In this case, genotypes with recent immigrant ancestry, that carry a *large* excess (i.e., 𝒪 (*L*)) of deleterious alleles relative to the average resident, exhibit essentially deterministic evolutionary dynamics. By contrast, more ‘average’ genotypes (that carry a *small* excess or deficit of deleterious alleles relative to the mean) will have dynamics that are significantly perturbed by drift.

Thus we have the following heuristic picture: a small subset of deleterious alleles embedded in immigrant or early-backcross genotypes experience strong negative selection due to statistical associations with other such alleles, but are largely unaffected by drift. Since such genotypes are rapidly broken down by recombination (at least under free recombination) and/or eliminated by strong selection, they contribute little to fitness variance in the recipient population. As a consequence, the *majority* of deleterious alleles are embedded in more average genotypes and are, thus, significantly affected by direct selection (due to their own deleterious effect) and genetic drift, but not indirect selection due to LD. Under these conditions, it is reasonable to expect that the equilibrium allele frequency distribution would be close to the single-locus distribution under drift-migration-selection-mutation balance, but with a *reduced* effective rate of migration, where the reduction reflects selection against immigrant and early-backcross genotypes, which causes sets of deleterious alleles to be eliminated together.

To make these arguments more concrete, consider first the mainland-island case. If the rate of drift 1*/N* is much higher than the (effective) rate of migration and mutation, allele frequency distributions will be U-shaped, and the island population near fixation for one or other allele at each locus. Denoting the expected deleterious allele frequency per locus under migration-selection-mutation-drift balance by 𝔼 [*p*], a U-shaped distribution implies that the island is near fixation for the locally deleterious allele at ≈ *L* 𝔼 [*p*] loci, so that migrants from the mainland carry an excess of *L*(1 − 𝔼 [*p*]) alleles on average. This suggests that the relevant reduced migration rate governing allele frequency distributions (and consequently the expected frequency 𝔼 [*p*]) can be approximated by *m*_*e*_[*s, L*(1 − 𝔼 [*p*])], where *m*_*e*_ is given by Eq (4).

We use *ψ*_*island*_ [*p* | 𝔼 [*p*]] to denote the allele frequency distribution, conditional on the expected allele frequency 𝔼 [*p*]. This is simply given by Eq (1), but with the raw migration rate *m* replaced by the effective migration rate *m*_*e*_[*s, L*(1 − 𝔼 [*p*])]. We can now obtain an implicit equation for 𝔼 [*p*] by using the fact that: 𝔼 [*p*]= *dp ψ*_*island*_ [*p* | 𝔼 [*p*]]. Thus, we have:

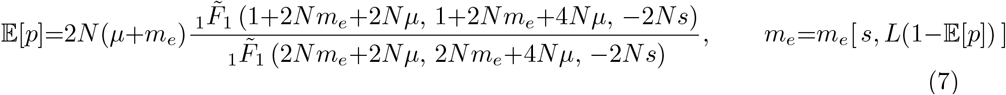

where 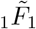 is the confluent hypergeometric function of the first kind. Eq (7) can be solved numerically to obtain 𝔼 [*p*].

Note that in replacing *m* (which is the migration rate *per generation*) by *m*_*e*_ (which is a composite parameter that encapsulates the splitting of the migrant genome over multiple, i.e., ∼ 10 generations of backcrossing), we implicitly assume that the splitting occurs much faster than changes due to any single-locus process, i.e., 1*/s*, 1*/m, N* « 1.

When allele frequency distributions at selected loci are not U-shaped and heterozygosity 𝔼 [*pq*] is appreciable, then the average fitness on the island is 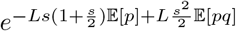. Thus, the relative selective disadvantage (and RV) of a migrant on the island will depend not only on the average number of selective differences between the mainland and island populations, but also on the heterozygosity (at selected loci) within the island population. However, the term involving heterozygosity is proportional to *s*^2^*L* and is thus much smaller (by a factor that is 𝒪 (*s*)) than the first term involving the expected allele frequency, provided individual selective effects are weak. Thus, we can neglect it to a first approximation.

Now consider the case of the infinite-island population with two habitats. If both habitats are locally adapted, then individuals migrating between demes within the same habitat will have higher RV, i.e., contribute more genetic material to future generations, than individuals migrating between demes belonging to different habitats. Thus, deleterious alleles can be associated with effective migration rates that now depend on both the habitat from which the allele originates as well as the habitat into which it immigrates.

Let 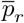 and 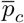 denote the expected frequencies in the rare and common habitat respectively (for the allele that is favoured in the common habitat). If drift is strong or at least comparable to other evolutionary processes, then any deme is close to fixation for one or other allele at each locus. In this scenario, immigrants will typically carry alleles that are positively selected vis-a-vis the resident allele at some loci and negatively selected at other loci. In principle, one can calculate the deterministic introgression dynamics of such a mosaic genome, containing both types of alleles [42]. However, as a first approximation, we will assume that all that matters is the net selective disadvantage of such a genome, which is, on average, proportional to the excess number of locally deleterious alleles that it carries, relative to a typical resident. This is 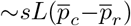 for individuals migrating *between* habitats, and zero for migrants *within* the same habitat.

As before, we can write down the distribution 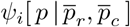 of allele frequencies on any island within habitat *i*, conditional on 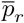 and 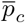 the average frequencies across the rare and common habitats respectively. Integrating over these yields 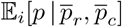, the expected frequency within any deme in habitat *i*, conditional on 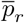 and 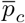. Finally, by using the fact that 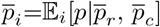 for all *i* (at equilibrium), we arrive at the following coupled equations for 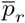 and 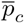:

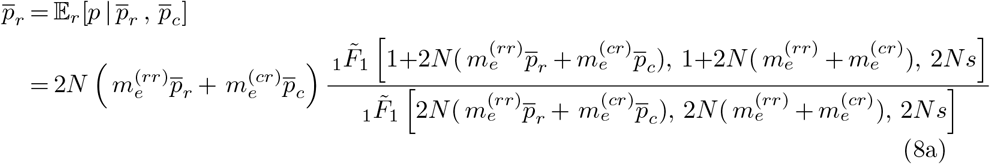

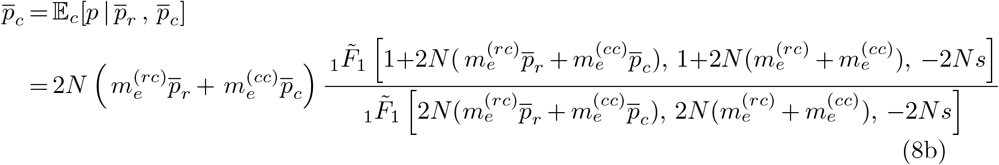

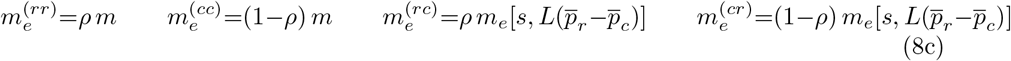

Thus, in this case, allele frequency distributions depend on an effective migration *matrix* with elements 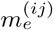, which denote the probabilities that a lineage sampled in habitat *j* in the present has originated from habitat *i* in the previous time step. For *i*=*j*, i.e., migration within the same habitat, 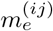 is assumed to be equal to the raw migration rate multiplied by the fraction of islands in the habitat of origin. For *i*≠*j*, i.e., migration between habitats, 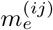 is approximated by the effective migration rate 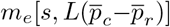 times the fraction of islands in the habitat of origin (see Eq (8c)). Eq (8) can be solved numerically to obtain 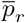 and 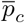 – the expected frequencies in the two habitats at equilibrium.

As in the mainland-island case, we neglect the contribution of heterozygosity (within demes) to effective migration rates, as this is much smaller than the contribution of the mean allele frequency difference 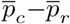 between demes. By the same token, in this first approximation, we can also neglect the contribution of the variance of the allele frequency difference (across different loci). As before, the contribution of these variances is smaller (by a factor that is 𝒪 (*s*)) than the contribution of the mean allele frequency difference.

### Barriers to gene flow and neutral divergence in the infinite-island model

Local adaptation elevates genomewide *F*_*ST*_ across all subpopulations by reducing the *effective* rate of immigration into any deme (since immigrants originating from the dissimilar habitat have low RV). However, effective immigration is more strongly reduced for demes in the rare than in the common habitat, since most immigrants into the former originate from a dissimilar habitat. Thus, we must consider habitat-specific statistics 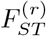 and 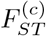 — these represent the probability of identity by descent (at an unlinked neutral locus) of two lineages sampled from a deme within the rare and common habitats respectively, relative to the probability of identity of two lineages sampled from anywhere within the entire population.

Following Slatkin (1991) [43], we can express the expected value of these F-statistics in terms of expected pairwise coalescence times: 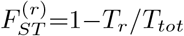 and 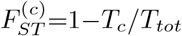, where *T*_*r*_, *T*_*c*_ and *T*_*tot*_ denote the average coalescence time of two lineages, both sampled from a single deme belonging to the rare habitat (*T*_*r*_), or both from a single deme within the common habitat (*T*_*c*_), or each lineage sampled independently from across the entire population (*T*_*tot*_).

Often, in practice, it is only possible to estimate neutral divergence between *pairs* of subpopulations, here labeled *i* and *j*. In this case, the expected 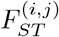 is: 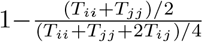, where *T*_*ii*_, *T*_*jj*_ and *T*_*ij*_ are, respectively, the expected coalescence times for pairs of lineages both sampled from deme *i*, or from deme *j*, or one from *i* and one from *j*. As before, we must separately consider the cases where both sampled demes belong to the rare habitat or to the common habitat or one to the rare and the other to the common habitat: this leads to three divergence measures 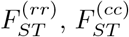 and 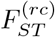

Let *m*_*rc*_ and *m*_*rr*_ denote the probability per unit time that a lineage sampled in the rare habitat traces back to the rare and common habitat respectively. We can define analogous backward migration rates *m*_*cc*_ and *m*_*cr*_ for lineages sampled in the common habitat. We can then obtain the various expected coalescence times using the structured coalescent (Section 2, S1 Text), and express these in terms of the population-size-scaled migration rates *M*_*rr*_=*Nm*_*rr*_, *M*_*rc*_=*Nm*_*rc*_, *M*_*cc*_=*Nm*_*cc*_, *M*_*cr*_=*Nm*_*cr*_. As before, the key approximation is to assume that *m*_*rc*_ and *m*_*cr*_ are attenuated by a factor that depends on the ‘barrier effect’ of the *L* selected loci, when the habitats are (partially) locally adapted (see also [19] for a similar approximation). More specifically, we will assume that *M*_*rc*_ and *M*_*cr*_ are proportional to 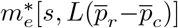, where 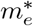 is given by Eq (5), and 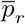 and 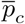 are obtained from Eq (8). Thus, we have (see Section 2, S1 Text for details):

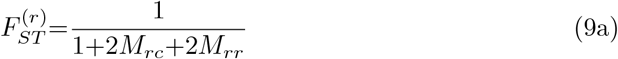

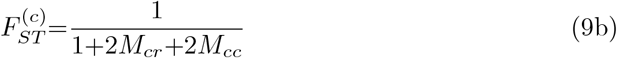

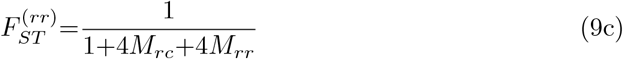

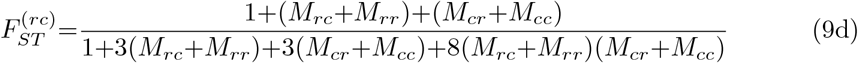

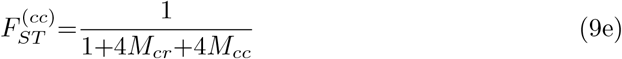

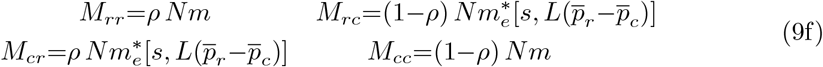

Note that Eq (9f) is similar to Eq (8c), but with the effective migration rate 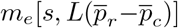 for selected alleles (Eq (4)) replaced by the corresponding rate 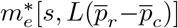 for neutral alleles (Eq (5)).

Since we assume equal and opposite selective effects of any allele in the two habitats (*s*_*c*_=*s*_*r*_=*s*), we have: (1− *ρ*)*M*_*rr*_=*ρM*_*cc*_ and (1−*ρ*)*M*_*cr*_=*ρM*_*rc*_ (Eq (9f)). From this, it also follows that 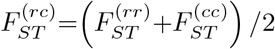. However, this does not hold generally, e.g., with asymmetric selection across habitats.

### Individual-based simulations

Individual-based simulations are used to check the accuracy of the various approximations introduced above. In case of the mainland-island model, a single focal deme with *N* haploid individuals carrying *L* selected unlinked loci is simulated. In each generation, a Poisson number of individuals with mean *Nm* is replaced by mainland individuals, who carry the locally deleterious allele at each of their *L* loci. Mutation is then introduced by flipping the state of each locus of each individual independently with probability *µ*. Selection is implemented by multinomial sampling of 2*N* parents (with replacement) from the existing *N* individuals by choosing sampling weights to be equal to (relative) individual fitness. *N* offspring in the next generation are then produced by randomly pairing parents and creating offspring genotypes by independently choosing the allelic state at each locus to be the same as that of either parent with equal probability.

The simulation procedure for the island model with *D* islands is the same (excluding mutation), except that migration is implemented by first removing a Poisson-distributed number of individuals (with mean *Nm*) from each deme to create a common migrant pool; individuals from this pool are then randomly redistributed back into the *D* demes, while ensuring that the size of each deme remains constant at *N*. Simulations are initialized such that there is maximum polymorphism (allele frequency 0.5) at each locus within each population; other initial conditions are considered in Section 6, S1 Text.

For the *D*-island model, an additional set of *L*_1_ neutral bi-allelic markers are simulated; these are unlinked to each other and to selected loci, and are also initialized with allele frequency 0.5. Once the population has equilibrated, the single-deme *F* statistics are calculated as: 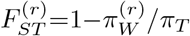 and 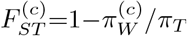, where 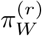 (respectively 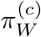) is the heterozygosity within demes in the rare (resp. common) habitat, averaged over all neutral markers and over all demes in the habitat; *π*_*T*_ is the diversity across the whole population, averaged over all neutral markers. Similarly, the 2-deme F measures are calculated as: 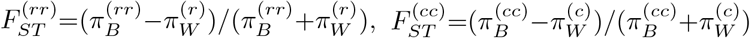 and 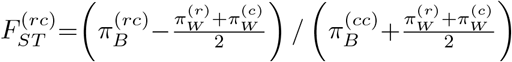. Here, 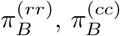 and 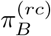 denote the number of pairwise differences per site between 2 demes belonging to the same or different habitats; these are calculated (as above) by averaging over all neutral markers and over all relevant pairs of demes.

## Data availability

A MATHEMATICA notebook for the numerical approximations and FORTRAN code for individual-based simulations is available at: https://github.com/HimaniSachdev/RI_LD_drift.

## Results

I first analyse the mainland-island model, focusing on how local adaptation on the island depends on *m/s* (the migration rate relative to per locus selection), *Ns* (the strength of selection per locus relative to drift), and *Ls* (the net selection against maximally deleterious genotypes; this is the main determinant of the magnitude of the barrier effect due to LD). Since the focus is on the interplay between migration, multi-locus selection and drift, the mutation rate is set to a fixed value throughout the main paper. The sensitivity of local adaptation thresholds to mutation is considered in Section 4, S1 Text.

I then consider the infinite-island model, where there is an additional parameter *ρ* (which parametrizes the relative abundances of the two habitats). I focus on the conditions under which LD widens the range of migration rates over which local adaptation can be maintained in the rare habitat, and on the effect of polygenic barriers on neutral divergence between habitats. Throughout, theoretical predictions (Eq (7-9)) using Eq (4-5) are compared against individual-based simulations.

### Mainland-island model

Fig 1A (main plot) shows the expected equilibrium frequency of the locally adaptive allele on the island as a function of *m/s*, for different numbers *L* of divergently selected loci (various colors), with the other parameters being *s*=0.02, *Ns*=2, *µ/s*=0.005. As expected, locally adaptive alleles become less common with increasing migration for any *L*. However, in contrast to the relatively smooth decrease in allele frequency with *m/s* observed for small *L*, there is a threshold effect when *L* is large— frequencies decrease only mildly with migration at low *m/s*, but then collapse beyond a critical migration threshold. Accordingly, adaptive allele frequencies can be much higher than the single-locus/LE prediction (shown in brown in Fig 1A), when *L* is large. This is simply a consequence of stronger multilocus selection against individuals with recent immigrant ancestry (at larger *Ls*), which causes sets of deleterious alleles to be eliminated together, before they can break up by backcrossing with fitter backgrounds.

The inset of Fig 1A shows the expected maladaptation load (calculated by summing across all loci) vs. *Ls*, where *Ls* is varied by varying *L*, while holding *s*=0.02 constant. The dashed line depicts the maximum possible load (also equal to *Ls* in this model): this corresponds to an extreme scenario where the island population is fixed for locally deleterious alleles at all loci. We see that load is actually highest for intermediate values of *Ls*, which can be understood as follows: at small *L*, any increase in the number of loci contributing to maladaptation far outweighs the milder decrease in deleterious allele frequency per locus. However, once *Ls* is large enough for selective elimination of groups of alleles to be effective, deleterious allele frequencies may decrease quite sharply with increasing *L*, which compensates for the increase in the number of loci, causing total load to decline with *L*.

Our focus on the composite parameter *Ls* is justified by the fact that effective migration rates depend primarily on *Ls*, rather than on *L* and *s* separately (see Eq (6)). The sensitivity of allele frequencies to the exact genetic basis of load, i.e., to *L* and *s* (for a given *Ls* and *Ns*) is explored in Section 3, S1 Text.

A striking feature of Fig 1A is the close agreement between the results of individual-based simulations (symbols) and theoretical predictions (lines), which are obtained by numerically iterating Eq (7). This rather successful approximation of the effects of LD on allele frequencies in terms of effective migration rates suggests the following explanation for sharp thresholds for loss of local adaptation at large *Ls*: a small increase in maladaptation at many loci may, in aggregate, cause a substantial increase in the effective migration rate *m*_*e*_ ≈ *me*^*−*2*Ls*(1*−* 𝔼 [*p*])^ associated with individual alleles, provided *Ls* is large. This, in turn, further increases swamping, pushing up frequencies of locally deleterious alleles, setting in motion a positive feedback, which culminates in collapse of local adaptation above a threshold migration rate.

Consider next how genetic drift influences local adaptation. Fig 1B shows the expected frequency of the locally favoured allele as a function of *m/s*, the migration rate relative to selection strength per locus, for island populations of different size (different colors). As before, simulation results (symbols) match theoretical predictions (colored solid curves) very well across all parameter combinations. Larger sizes (which correspond to higher values of *Ns*) allow populations to sustain local adaptation at significantly higher migration levels. For example, while the frequency of the locally adaptive allele drops below 0.1 already at *m/s*≈0.25 in the smallest population (with *Ns*=1), the corresponding threshold is *m/s*≈0.98 in the largest population (with *Ns*=16), close to the deterministic threshold *m/s*≈1.11. Here, the deterministic equilibrium allele frequency *p*_*det*_ at migration-selection-mutation balance is obtained by numerically solving: −*s p*_*det*_(1−*p*_*det*_)+*m*_*e*_[*s, L*(1−*p*_*det*_)](1−*p*_*det*_)+*µ*(1−2*p*_*det*_)=0, and using the approximate expression in Eq (6a) for *m*_*e*_. This prediction (shown via the solid black curve in Fig 1B) accounts for the effects of LD between introgressing deleterious alleles but neglects drift. A comparison of the finite *Ns* plots with the deterministic prediction shows that genetic drift has a significant effect on local adaptation for *Ns ≲* 10 in this example.

Fig 1B also shows LE/single-locus predictions for each *Ns* (colored dashed curves; obtained from Eq (1)): these account for the effects of genetic drift but neglect LD. The LE prediction for the deterministic allele frequency *p*_*det,LE*_, obtained by solving −*s p*_*det,LE*_ (1 − *p*_*det,LE*_)+*m*(1 − *p*_*det,LE*_)+*µ*(1 − 2*p*_*det,LE*_)=0, is also shown for reference (black dashed curve). While the frequency of the locally favoured allele (symbols/solid lines) is higher than the corresponding LE prediction (dashed lines) at all *Ns*, this effect is stronger in larger populations, which also exhibit sharper thresholds for loss of local adaptation. This can be understood, as before, in terms of the effect of genetic drift on effective migration rates— larger populations are less likely to fix locally deleterious alleles and thus have lower load. Consequently, migrants from the mainland have lower relative fitness in these populations (all other parameters being equal), causing effective immigration rates to be also lower in larger populations, thus further protecting locally adaptive alleles from swamping.

At first glance, it is surprising that this simple heuristic based on introducing effective migration rates into the single-locus diffusion approximation should accurately predict the expected allele frequency across such a wide range of parameters, including in large populations, where one or other allele is not necessarily close to fixation. This behoves us to ask: how sensitive are these approximations to assumptions about the (U-)shape of the underlying allele frequency distribution? We can investigate this by contrasting frequency distributions in populations of different size (Fig 1C), choosing the migration rate in each case such that the expected deleterious allele frequency is ∼ 0.3 (based on the theoretical prediction), regardless of size.

As expected, the allele frequency distribution is U-shaped in smaller populations, but unimodal and peaked around 𝔼[*p*] ∼ 0.3 in larger populations. Interestingly, theoretical predictions for the frequency distribution (lines) are quite accurate even in the two largest populations (purple and blue triangles in Fig 1C, corresponding respectively to *Nm* ∼ 6 and *Nm* ∼ 13), for which distributions of the selected allele deviate markedly from the canonical U-shape, and are characterised by high heterozygosity (𝔼[2*pq*] ∼ 0.365 and ∼ 0.399 respectively). Thus, this rather crude representation of multi-locus LD via a single effective migration rate, that depends only on the expected number of genetic differences between the mainland and island, appears to suffice even if there is substantial polymorphism, i.e., if these differences represent pairwise differences between individuals rather than fixed differences between populations. This is likely due to the fact that, to lowest order in *s*, the effective migration rate only depends on the average divergence between and not heterozygosity within populations (see Model and Methods).

Finally, we ask: can the approximations introduced here predict the equilibrium *genotype* frequency distribution? Fig 1D shows the equilibrium frequencies *P*_*y*_ for genotypes carrying *y*=0, 1, 2, … deleterious alleles, for two values of *m*, as found in simulations (symbols), along with two kinds of analytical predictions (solid and dashed lines). The solid lines show the predicted genotype frequencies *under LE* given the expected deleterious allele frequency 𝔼 [*p*] (which is determined as described above, using Eq (7)). Under LE, the allelic states at different loci are statistically uncorrelated, and the probability of genotypes with exactly *y* deleterious alleles is 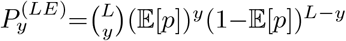.

We also compare against the *deterministic* prediction for *P*_*y*_ (dashed lines in Fig 1D) obtained by solving Eq (3) (see also eq. 1 in S1 Text): more specifically, the dashed lines show 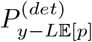 vs. *y*, where the deterministic frequencies *P* ^(*det*)^ are calculated by assuming that the island is subject to divergent selection at *L*(1− 𝔼 [*p*]) loci. Note that the relevant deterministic frequencies are 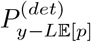 and not 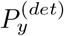 as genotypes carrying *y* deleterious alleles have relative (Malthusian) fitness that is proportional to *y* − *L* 𝔼 [*p*] (since the island is nearly fixed for *L* 𝔼 [*p*] deleterious alleles, by definition).

At the lower migration level (red plot in Fig 1D), genotype frequencies *P*_*y*_ are close to the LE prediction (solid line) for small *y*, while they match the deterministic prediction (dashed line) for large *y*, with a crossover between the LE and deterministic regimes at intermediate *y*. This suggests that when migration is low and deleterious alleles correspondingly rare, the frequencies of genotypes with large numbers of deleterious alleles (large *y*) are governed by the (deterministic) balance between migration, selection and recombination, with recombination only serving to break down highly deleterious genotypes into smaller, less deleterious fragments, but rarely bringing together such fragments to reconstitute the more deleterious genotypes (this is tantamount to assuming that { *P*_*y*_ } satisfy *linear* coupled equations, as in Eq (3)). In contrast, both roles of recombination— the splitting of more deleterious genotypes to generate the focal genotype and the reconstitution of the focal genotype via recombination between less deleterious genotypes, appear to play a role in shaping the frequencies of genotypes with low numbers of deleterious alleles (small *y*): this is reflected in the fact that *P*_*y*_ for small *y* are close to the LE prediction, which assumes that genotypes are random assortments of independently segregating alleles. We note that *P*_*y*_ are slightly elevated above the LE prediction for very small *y*, reflecting positive selection on such genotypes due to their higher relative fitness.

At the higher migration level (blue plot), the distribution does not exhibit two distinct regimes. In particular, at large *y*, the *P*_*y*_ are much higher than the deterministic prediction, suggesting that when migration is high and deleterious genotypes more common, mating events which bring together different sets of deleterious alleles to generate highly deleterious genotypes are also relevant; accounting for such events would introduce terms of the kind *P*_*y*_*P*_*y*_*′* in Eq (3)). Also, the actual distribution of *P*_*y*_ for smaller *y* is significantly wider than the LE prediction in this case, suggesting significant selection not just on individual alleles but also sets of alleles.

The fact that these approximations nevertheless accurately predict allele frequency distributions at high levels of migration suggests that the heuristic of effective migration rates captures the *gross* effects of LD on allele frequencies quite robustly, even when it does not provide a good handle on LD (or equivalently, on genotype frequencies) itself. As before, this is a reflection of the fact that effective migration rates are insensitive to within-deme genetic variance (to lowest order in *s*), and thus, are relatively insensitive to within-deme LD (since it only contributes to the variance within demes).

### Infinite-island model

Let us now consider adaptive divergence between habitats in the infinite-island setting, where a non-zero fraction *ρ* of islands belong to the rare habitat. In this case, both habitats influence each other via maladaptive gene flow. Thus, we must consider how allele frequencies in the two habitats co-evolve, instead of assuming the state of the more abundant habitat to be ‘fixed’ and independent of the rare habitat, as in the mainland-island case.

In the absence of mutation, there is a well-defined critical migration rate *m*_*c*_, such that locally adaptive alleles can segregate in the rare habitat for *m<m*_*c*_, but are eliminated for *m>m*_*c*_. In other words, above this threshold, no long-term adaptive divergence between habitats is possible— instead, alleles favoured in the common habitat fix across all islands, irrespective of habitat. Before analysing the effects of drift and LD on this critical threshold systematically, it is useful to consider a few examples.

Figs 2A and 2B show the expected frequency of the locally adaptive allele in the rare habitat, as a function of *m/s*, the migration rate relative to per-locus selection, for different *Ns* (blue vs. black colors), different *L* (different symbols in each subfigure), and different *ρ* (right vs. left columns). As before, theoretical predictions that account for both drift and LD (solid lines), obtained by numerically solving Eq (8) in conjunction with Eq (4), are in good agreement with the results of individual-based simulations (symbols) across all parameter combinations. LE/single-locus predictions that account for drift (but not LD) are also shown for reference (dashed lines).

**Fig 2.**
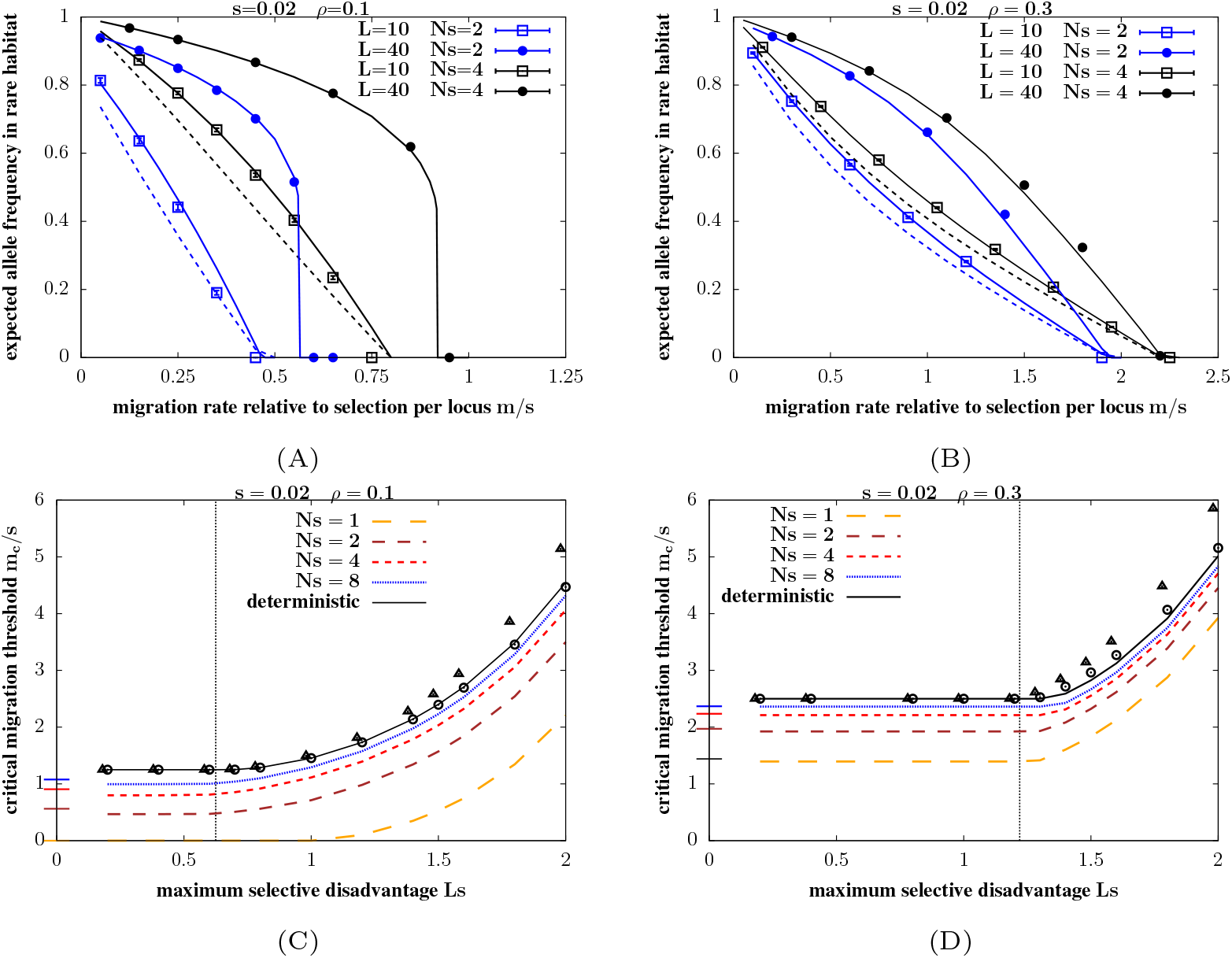
Local adaptation in the infinite-island model with two habitats. A–B. Expected equilibrium frequency of the locally adaptive allele in the rare habitat vs. *m/s* for *ρ*=0.1 (Fig A) and *ρ*=0.3 (Fig B), for 2 different values of *L* (10 and 40; squares vs. circles), and two different population sizes (corresponding to *Ns*=2 and *Ns*=4; blue vs. black). Symbols depict results of individual-based simulations; solid lines depict theoretical predictions that account for both LD and drift (obtained using Eq (8) together with Eq (4)); dashed lines depict LE (i.e., single-locus) predictions that only account for drift (obtained from Eq (2)). Selective effect per deleterious allele is *s*=0.02 in both plots. The number of simulated demes is *D*=500 in all individual-based simulations; the average allele frequency is obtained by averaging over all *L* loci and all *ρ D* islands in the rare habitat, across 5 simulation replicates. C–D. Theoretical predictions for *m*_*c*_*/s*, the critical migration threshold scaled by the per-locus selection coefficient, vs. *Ls* for *ρ*=0.1 (Fig C.) and *ρ*=0.3 (Fig D.), for *s*=0.02 and *N* =50, 100, 200, 400 (corresponding to *Ns*=1, 2, 4, 8 respectively). Here, *m*_*c*_ is the critical migration threshold above which local adaptation cannot be maintained in the rare habitat. Theoretical predictions are obtained by solving for the polymorphic equilibrium of Eq (8) (using Eq (4)) and determining the value of *m* above which no such equilibrium exists. The short horizontal colored lines along the vertical axis represent the approximate LE (single-locus) prediction 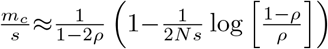 (see [38]). The exact deterministic predictions for *m*_*c*_*/s* (obtained by solving coupled deterministic equations for *p*_*r*_ and *p*_*c*_; see eq. 12 in Section 5, S1 Text) are shown using solid black lines. The critical migration rate *m*_*c*_*/s* is constant for small *Ls*, but then starts increasing with *Ls* beyond a threshold (*Ls*)_*∗*_. The deterministic prediction for (*Ls*)_*∗*_ is depicted by vertical dotted lines and depends only on the habitat fraction *ρ* (see text). In addition, we also show approximate deterministic predictions for *m*_*c*_*/s* (triangles and circles)— for *Ls>*(*Ls*)_*∗*_ and in the highly polygenic limit *s* → 0, *L* → ∞ with *Ls* constant, the deterministic *m*_*c*_*/s* is given by Eq (10b) and is shown using triangles. Predictions that are more accurate atl somewhat larger *s* are obtained in Section 5, S1 Text (see eq. 17B in S1 Text). These are shown using circles and agree well with the numerically obtained deterministic *m*_*c*_*/s* (solid black lines).

Comparing Figs 2A and 2B, we see that the critical migration threshold is higher for larger *ρ*: thus, local adaptation in the rare habitat can be sustained over a wider range of migration rates when the relative abundances of the two habitats are more similar (so that a smaller fraction of genotypes immigrating into the rare habitat are deleterious). Further, *m*_*c*_*/s* also increases with *Ns*, especially for *ρ*=0.1 (blue vs. black plots in Fig 2A), suggesting that drift has a significant effect on local adaptation in rare habitats in this parameter regime.

For the parameters depicted here, the critical threshold is approximately the same for *L*=10 and *L*=40, and thus is insensitive to *Ls*, the total selection difference between habitats. As discussed below, this is only true if *Ls* is lower than a threshold (*Ls*)_*∗*_, which depends on *ρ*. For *Ls>*(*Ls*)_*∗*_, LD between locally adaptive alleles allows adaptation to be maintained over a much larger range of migration rates than would be possible for selection acting on one locus alone (see Figs 2C and 2D).

Note that higher values of *Ls* always result in *sharper* thresholds for loss of adaptation across all parameter combinations, even when the critical migration rate *m*_*c*_ (at which adaptation collapses) remains unchanged. As before, sharper thresholds are a consequence of stronger LD between loci at higher *Ls*. Thresholds are also sharper for smaller *ρ*, which can be rationalised as follows: sharp thresholds emerge when an increase in migration causes a substantial enough increase in load in the recipient population that the effective migration rate of deleterious alleles immigrating from the *alternative* habitat also rises significantly (due to a rise in the RV of migrants between habitats), setting in motion a positive feedback between declining population fitness and rising (effective) maladaptive immigration into the population. Note that this kind of positive feedback only involves alleles migrating between differently adapted habitats, and not deleterious alleles that migrate within the same habitat (see also Eq (8c)). Thus, feedback effects are stronger and the threshold for loss of local adaptation on an island within the rare habitat sharper if immigration into the island is predominantly from the alternative habitat (as is the case if *ρ* is small).

When does LD cause the critical migration threshold for loss of local adaptation to *shift* (in addition to becoming sharper), and to what extent are such shifts opposed by genetic drift? One can investigate this question systematically by plotting *m*_*c*_*/s* vs. *Ls* for various values of *N* (or equivalently, *Ns*, since *s*=0.02 is held constant). Here, *m*_*c*_ is the critical migration threshold above which no adaptive divergence is possible, regardless of the initial state of the population, *Ls* is a proxy for the (maximum possible) barrier effect due to LD, and 1*/*(*Ns*) measures the strength of genetic drift relative to selection per locus. Figs 2C and 2D show theoretical predictions for *m*_*c*_*/s* for various *Ns* (colored dashed lines) for *ρ*=0.1 and *ρ*=0.3 respectively. To disentangle the effects of LD and drift, it is useful to also consider the deterministic (*N* →∞) predictions for *m*_*c*_*/s* (shown by solid black lines). Details of the deterministic analysis are presented in Section 5, S1 Text; only the main findings are summarised here.

### Effect of LD on critical migration thresholds (neglecting drift)

The deterministic analysis identifies a threshold (*Ls*)_*∗*_, above which LD between locally adaptive alleles is strong enough to raise the critical migration rate. The threshold (*Ls*)_*∗*_ depends only on the relative abundances of the two habitats (in the deterministic limit), and is given by (*Ls*)_*∗*_ ≈ 1*/*[2(1 − 2*ρ*)] for *ρ* ≤ 1*/*4, and (*Ls*)_*∗*_ ≈ 4*ρ* for 1*/*4*<ρ<*1*/*2 (see Section 5, S1 Text). These thresholds are indicated by dotted vertical lines in Figs 2C and 2D.

For *Ls<*(*Ls*)_*∗*_, there exists a single critical migration threshold, *m*_*c*,1_ */s*=1*/*(1−2*ρ*), which is independent of *Ls*. Both habitats evolve (partial) local adaptation for *m<m*_*c*,1_, resulting in non-zero allele frequency divergence between habitats at equilibrium. Conversely, no divergence can be maintained when *m>m*_*c*,1_, regardless of initial levels of divergence. Note that *m*_*c*,1_is simply the threshold for the maintenance of polymorphism at deterministic migration-selection equilibrium for a *single* locus.

For *Ls>*(*Ls*)_*∗*_, we observe two thresholds *m*_*c*,1_(which is the LE/single-locus threshold for polymorphism, discussed above) and a second threshold *m*_*c*,2_*>m*_*c*,1_(described below). There is always stable adaptive divergence between habitats at low migration rates, i.e., for *m<m*_*c*,1_, while migration necessarily erodes all adaptive divergence for *m>m*_*c*,2_(even when the two habitats are initially perfectly locally adapted). At intermediate migration rates, i.e., for *m*_*c*,1_*<m<m*_*c*,2_, evolutionary outcomes depend on the *initial* state of the metapopulation. In particular, long-term divergence is possible either if initial divergence between habitats is high (as in a scenario of secondary contact between subpopulations that have diverged in allopatry) or if the rare habitat harbours sufficient adaptive variation that multilocus divergence can build up and LD-mediated barrier effects emerge faster than migration washes out allele frequency differences at individual loci.

Note that the rising part of the black curves in Figs 2C and 2D corresponds to *m*_*c*,2_since the plots show critical migration thresholds above which long-term divergence is not possible, regardless of the initial state of the population.

The deterministic predictions for *m*_*c*_*/s* shown in Figs 2C and 2D (solid black lines) are obtained by numerically solving the deterministic equations for *p*_*c*_ and *p*_*r*_ (see eq. 13 in Section 5, S1 Text). However, an approximate analytical expression for *m*_*c*,2_can be obtained in the limit of highly polygenic divergence (i.e., for *s* → 0, *L* → ∞ with *Ls* constant). This expression is given by Eq (10a) below and is shown using triangles in Figs 2C and 2D. Moreover, in this limit, we can also obtain the critical allele frequency divergence Δ_*c*_=*p*_*c*_−*p*_*r*_ between habitats, as *m* approaches *m*_*c*,2_(see Eq (10b)).

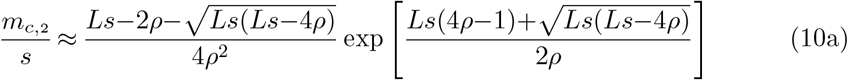

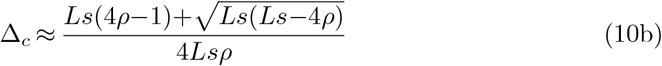

Thus, allele frequency divergence between habitats must be at least Δ_*c*_ for LD between adaptive alleles to maintain local adaptation. Once divergence falls below this level, there is a positive feedback between an increase in maladaptation load in the rare habitat and a corresponding increase in effective migration rate of deleterious alleles from the common to the rare habitat, leading to a sharp collapse in adaptive divergence.

### Effect of drift (and LD) on critical migration thresholds

Let us now consider critical migration thresholds in populations where *Ns* is not too large and drift has significant effects on single-locus polymorphism. These thresholds are depicted by the various colored dashed curves in Figs 2C and 2D, and are obtained by numerically finding the migration rate above which no polymorphic fixed point (with 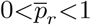) of Eq (8) exists.

As in the deterministic limit, *m*_*c*_*/s* is independent of *Ls* for small *Ls*, and is equal to the single-locus (LE) threshold, which is approximately: 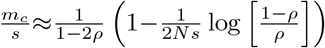 for *ρ* not too small [38]. These single-locus predictions (with drift) are indicated by horizontal colored dashes along the vertical axis in Figs 2C and 2D. Thus, in this regime, drift reduces the critical migration threshold *m*_*c*_*/s* by an amount proportional to 1*/*(*Ns*), with the reduction being more significant when one habitat is much rarer than the other (i.e., for smaller *ρ*). In fact, adaptive alleles are necessarily lost from the rare habitat in the absence of LD-mediated effects if 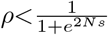, or equivalently, 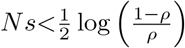, *regardless* of migration level (see e.g., *Ns*=1 curve in Fig 2C).

As before, sufficiently strong LD may raise the critical migration rate *m*_*c*_*/s* above the single-locus threshold. This requires *Ls>*(*Ls*)_*∗*_, where the threshold (*Ls*)_*∗*_ appears to be insensitive to *Ns*, and is thus predicted by the deterministic analysis (dashed vertical lines in Figs 2C and 2D; see also above). Interestingly, even with 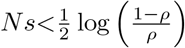, i.e., when selection on individual alleles is too weak to counter *any* level of gene flow in the long run, LD between many such alleles can maintain local adaptation under moderate levels of migration— this is, for example, the case for *Ns*=1 in Fig 2C.

Note that the rising part of the *m*_*c*_*/s* vs. *Ls* curves has approximately the same shape in populations with large *Ns* (e.g., blue and red dashed curves) as in the deterministic case (black solid curve), and is merely shifted downwards with respect to the deterministic prediction. However, at smaller *Ns*, the shape is also affected, with *m*_*c*_*/s* rising more slowly with *Ls* than in larger populations. These effects are, however, rather modest, suggesting that the main effect of drift is to cause a fixed reduction (which is more significant for smaller *Ns* and smaller *ρ*) in the critical migration threshold *m*_*c*_*/s*, regardless of *Ls*.

As in the deterministic case, whether or not polygenic local adaptation can evolve despite high levels of migration (via the emergence of strong multilocus barriers to gene flow) depends on the initial level of adaptive variation available, especially in the rare habitat. Section 6, S1 Text explores the behaviour of individual-based simulations initialized with different allele frequencies, for a few representative parameter combinations. With low initial adaptive variation in the rare habitat, populations may not be able to evolve enough divergence for LD-mediated effects to come into play. Then critical migration thresholds are lower than those depicted in the rising part of the *m*_*c*_*/s* vs. *Ls* curves in Figs 2C and 2D (though they may still be higher than the corresponding single-locus threshold, depending on the level of initial variation; see Section 6, S1 Text).

### Barriers to gene flow and neutral divergence

Let us now consider how adaptive divergence influences genetic diversity at unlinked *neutral* loci. In the absence of such divergence, *F*_*ST*_ depends only on *Nm*, the average number of migrants exchanged per generation between any island and the full population. Here, *Nm* may be viewed as a proxy for the physical subdivision of the population, which affects all subpopulations equally (under the island model). However, if habitats are locally adapted, then *F*_*ST*_ depends not only on the number of migrants exchanged but also on the RV of migrants, i.e., their long-term contribution to the neutral gene pool of the receiving population. As shown in this section (Fig 3), this dependency is captured accurately by expressing *F*_*ST*_ in terms of the elements of an effective migration rate matrix for neutral alleles (Eq (9)) and approximating effective migration rates as in Eq (5).

**Fig 3.**
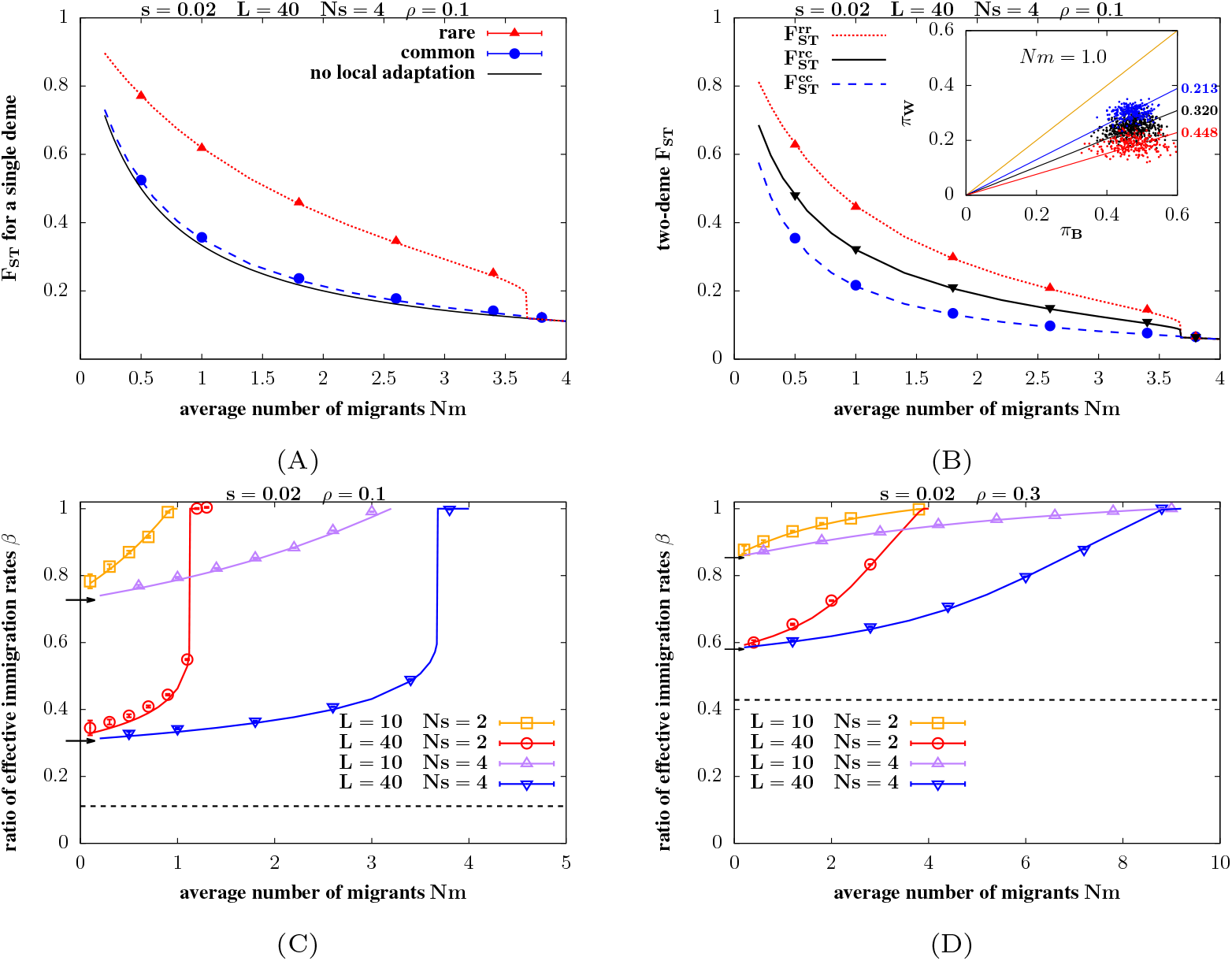
Neutral divergence in the infinite-island model. A. Average *F*_*ST*_ for a *single* deme in the rare 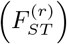 and common 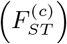 habitats vs. *Nm*, for *s*=0.02, *L*=40, *Ns*=4, *ρ*=0.1. Here, *F*_*ST*_ is measured relative to the whole metapopulation. Symbols show results of individual-based simulations; dashed lines represent theoretical predictions (obtained from Eqs (9a),(9b),(9f) and using Eq (5)); the solid black line represents *F*_*ST*_ =1*/*(1+2*Nm*)– the prediction in the absence of local adaptation. B. Average *F*_*ST*_ for a *pair* of demes vs. *Nm*, for the same parameters as in Fig A. Here, both demes within the pair may belong to the rare habitat 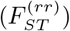, or both to the common habitat 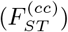, or one to the rare and the other to the common habitat 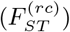. The plots show simulation results (symbols) as well as theoretical predictions (lines; obtained from Eq (9c)-(9f) and using Eq (5)). *Inset* (Fig 3B): The neutral diversity within demes *π*_*W*_ vs. neutral divergence between demes *π*_*B*_ for rare/rare, common/common and rare/common pairs of demes (shown using red, blue and black points respectively), for *Nm*=1.0, as measured at a single timepoint in an individual-based simulation. Each point represents a pair of demes (*i, j*); *π*_*W*_ is computed as 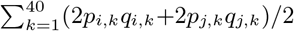 and *π*_*B*_ as 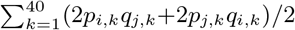, where *p*_*i,k*_ represents the allele frequency at the *k*_*th*_ neutral locus in deme *i* and *q*_*i,k*_=1*−p*_*i,k*_. The solid lines represent (*π*_*W*_, *π*_*B*_) combinations that would correspond to *F*_*ST*_ values of 0 (orange), 0.213 (blue), 0.320 (black), 0.448 (red): the last three are the predicted 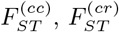 and 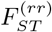 at *Nm*=1 respectively. C– D. 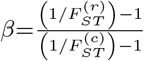 vs. *Nm* for *ρ*=0.1 (Fig. C) and *ρ*=0.3 (Fig. D), for different values of *L* and *Ns* (depicted by the different colors) and *s*=0.02. The quantity *β* gives the ratio of the *effective* number of immigrants per unit time into an island in the rare habitat to the corresponding number for an island in the common habitat. Theoretical predictions (solid lines) match simulation results (symbols) across all parameter combinations. The black dashed line in each plot represents the threshold *β*_*min*_=*ρ/*(1*−ρ*), which is the expected *β* under complete RI (wherein immigrants from the dissimilar habitat have zero RV). The short horizontal arrows along the vertical axis represent the threshold *β*=(*β*_*min*_+ *e*^*−*2*Ls*^)*/*(1+*β*_*min*_ *e*^*−*2*Ls*^) for *L*=10 (upper arrow) and *L*=40 (lower arrow). This is the expected value of *β* when allele frequency divergence between habitats is maximum (see text). *F*_*ST*_ values in simulations are computed from 40 unlinked neutral loci.

The rationale for focusing on *unlinked* neutral markers is that reduced gene flow at such markers signals the emergence of genome-wide, as opposed to localized, barriers to gene flow, and is thus a more appropriate measure of RI between habitats. Moreover, the barrier effect at *any* neutral site (even those in the vicinity of a particular barrier locus) is primarily due to unlinked loci, as long as selection acts on very many alleles of weak effect spread across a long genetic map [13].

Fig 3A shows average 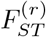 and 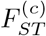 for islands belonging respectively to the rare and common habitat, as a function of *Nm*. The expected *F*_*ST*_ in the absence of local adaptation, which is 1*/*(1+2*Nm*) for haploids, is also shown (solid black line). As before, theoretical predictions (lines, obtained from Eqs (9a), (9b), (9f), in conjunction with Eq (5)) are in close agreement with simulations (symbols). Adaptive divergence between habitats increases neutral *F*_*ST*_ in both habitats above the neutral expectation (solid line), but more so in the rare habitat. This can be understood by noting that 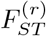 and 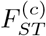 measure respectively the extent to which neutral diversity within any deme belonging to the rare or common habitat is reduced relative to gene diversity at the level of the population as a whole; this in turn depends on the effective rate of immigration into the deme. When both habitats are locally adapted, the majority (i.e., a fraction 1 − *ρ*) of immigrants into the rare habitat have very low RV, as they originate from the common habitat and thus carry genotypes that are locally deleterious. By contrast, only a minority (i.e., a fraction *ρ*) of immigrants into the common habitat have low RV. This results in a stronger reduction in neutral diversity and concomitantly, a sharper increase in *F*_*ST*_ for islands in the rare as compared to the common habitat.

We can also measure *F*_*ST*_ between pairs of islands in the population. Fig 3B (main plot) shows theoretical predictions (lines) and simulation results (symbols) for average 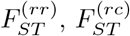 and 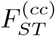, which correspond respectively to two islands both belonging to the rare habitat, or one to the rare and the other to the common habitat, or both to the common habitat. Note that 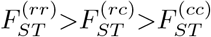: thus, at low *Nm, F*_*ST*_ between two islands in the rare habitat is *greater* than *F*_*ST*_ between islands from different habitats, despite the fact that only the latter exhibit adaptive divergence.

This somewhat paradoxical observation is explained by the fact that the rare habitat, though rare, is nevertheless large enough to harbour high genetic diversity (at the level of the habitat as a whole), as long as it encompasses a finite fraction of all islands in a large metapopulation. Thus, any two islands within the rare habitat exhibit high neutral divergence with respect to each other (since they will each be close to fixing a random set of alleles segregating in the habitat as a whole), even as the diversity within each is strongly reduced because of very low net effective immigration. By contrast, islands that support the common habitat have fairly high within-island diversity as well as between-island divergence, causing *F*_*ST*_ to be lower for any pairwise comparison that includes such islands. This is illustrated in the inset of Fig 3B by plotting neutral diversity within demes, *π*_*W*_, vs. neutral divergence between demes, *π*_*B*_, for *Nm*=1, for pairs of demes belonging to the same vs. different habitats (different colors). This plot shows that differences in *F*_*ST*_ are purely due to differences in *π*_*W*_, as *π*_*B*_ is the same for any pair of demes (on average), irrespective of habitat.

Thus, different levels of *F*_*ST*_ for islands within the rare vs. common habitat reflect the underlying asymmetry in the effective levels of immigration into the two kinds of islands when both are at least partially locally adapted. This asymmetry, in turn, is due to the fact that the majority of immigrants into the rare habitat have low RV, while this is true of only a minority of immigrants into the common habitat. This asymmetry can be measured directly via 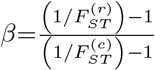, which is the ratio of the effective number of immigrants per generation into islands within the rare habitat to the corresponding number for islands in the common habitat.

In the absence of local adaptation (i.e., if alleles favoured in the common habitat are fixed across the entire population at all divergently selected loci), the relative fitness of all immigrants between and within habitats is equal to that of residents for any island. Then, the effective number of immigrants into any island is equal to the actual number, irrespective of habitat (at least under soft selection), so that *β*=1. In the opposite limit of complete RI, individuals migrating between habitats have RV close to zero, while migrants within a habitat have RV close to 1. Then, the effective number of immigrants into any island is the same as the number of immigrants that originate from other islands within the same habitat, so that 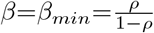.

Figs 3C and 3D show *β* vs. *Nm* for *ρ*=0.1 (Fig 3C) and *ρ*=0.3 (Fig 3D), for various values of *Ns* and *L* (same as those in Fig 2). As before, symbols represent results of individual-based simulations, while lines represent theoretical predictions; the horizontal dashed lines represent *β*_*min*_, which is the expectation under complete RI. As expected, *β* decreases as *Nm* decreases due to the concomitant increase in adaptive divergence between habitats. However, *β* always lies above the threshold *β*_*min*_ (dashed line) even for the smallest *Nm* that we simulate, indicating that RI is *incomplete*, even at very low levels of migration, as long as *Ls* is not very large. More precisely, we have: 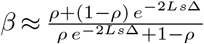, where 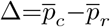 is the allele frequency divergence between two habitats, and *e*^*−*2*sL*Δ^ the RV of migrants between habitats. Thus, even with complete allele frequency divergence (Δ ≈ 1), *β* will be only be (*β*_*min*_+ *e*^*−*2*Ls*^)*/*(1+*β*_*min*_ *e*^*−*2*Ls*^), which is depicted by short horizontal lines along the vertical axis in Figs 3C and 3D.

Note that the typical values of *F*_*ST*_ found here— 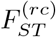 in the range 0.2−0.6 for *Nm<*2 (Fig 3B), correspond to the so-called “grey zone” of speciation, where it may be difficult to estimate effective migration rates and the extent of RI from sequence data using standard demographic inference methods [44]. The analysis here suggests that such values of *F*_*ST*_ are consistent with incomplete RI (and thus low levels of ongoing gene flow between habitats), at least in the context of patchy populations.

## Discussion

This paper introduces a simple heuristic for approximating the combined effects of LD and genetic drift on allele frequencies, when multiple loci are under divergent selection across distinct habitats in a subdivided population. It thus extends previous theory on polygenic barriers to gene flow (e.g., [5], [13]) to account for the effects of genetic drift within sub-populations. Drift may be significant when barrier loci have modest effects individually and/or when the scale of density regulation in populations is sufficiently local that *Ns* is small. However, divergent selection involving many such loci (large *Ls*) may still allow for local adaptation and substantial genomewide divergence between habitats, if migration levels are below a critical threshold.

### Effect of LD on critical migration thresholds for loss of local adaptation

A key finding of this study is that LD between adaptive alleles increases the critical migration threshold for loss of local adaptation only if *Ls* (which governs the strength of divergent selection) is above a threshold (*Ls*)_*∗*_. This threshold depends primarily on the relative proportions of the two habitats in the population (neglecting drift): for instance, when the rare habitat encompasses 20% of demes (*ρ*=0.2), we have (*Ls*)_*∗*_ ≈ 0.83, which corresponds to a relative immigrant fitness of 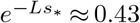 under conditions of very low gene flow (i.e., when alternative alleles are fixed at all *L* loci across habitats).

Below this threshold, i.e., for *Ls<*(*Ls*)_*∗*_, LD is not strong enough to maintain local adaptation beyond the single-locus migration threshold *m*_*c*,1_, though it does increase the extent of adaptive divergence between habitats for *m<m*_*c*,1_. By contrast, with *Ls>*(*Ls*)_*∗*_, LD between alleles can maintain adaptive divergence at migration levels that may be several times higher than the *m*_*c*,1_(see Figs 2C and 2D). The protective effects of LD on local adaptation are especially marked in very rare habitats, allowing these to withstand comparable levels of maladaptive migration as habitats that encompass a larger fraction (though still a minority) of islands. For example, in the absence of LD, the deterministic single-locus threshold for maintenance of adaptation in the rare habitat (*m*_*c*,1_*/s*=1*/*(1 − 2*ρ*)) is 1.25 for *ρ*=0.1 and 2.5 for *ρ*=0.3 (i.e., greater by a factor of 2 for the larger *ρ* value). However, at *Ls*=1.5, i.e., with LD, the corresponding deterministic thresholds (given by Eq (10b); see also Section 5, S1 Text) are ≈2.6 for *ρ*=0.1 and ≈3.2 for *ρ*=0.3, (i.e., greater by a factor of only 1.2).

An important caveat of this analysis is that we assume that there is always enough standing genetic variation to allow for a rapid buildup of initial divergence across habitats (though see Section 6, S1 Text for some examples with limited initial variation). This is then further reinforced by the positive feedback between increasing divergence and falling effective migration rates, provided *Ls>*(*Ls*)_*∗*_ and migration is below a certain threshold. While rapid adaptation from standing genetic variation (possibly within a hybrid swarm) has been implicated in adaptive radiations [45, 46], this is hardly the norm. More generally, if adaptive divergence is mutation-limited to at least some degree, then initial divergence would build up slowly and may be seeded by pre-existing adaptive differences (so-called divergence hitchhiking [14]), until net divergence reaches a high enough value that genomewide effective migration rates drop, causing genomes in the different populations to ‘congeal’ [21, 22].

Here, we identify a related threshold: if allele frequency divergence falls below Δ_*c*_, there is a sharp increase in the genomewide effective migration rate, and divergence collapses. In the previous example with *ρ*=0.2 and assuming *Ls*=1, allele frequency divergence between habitats must be at least ≈0.3, i.e., migrant fitness no greater than 0.73 relative to population mean fitness and *m*_*e*_*/m* no greater than 0.54, for congealing to be possible, and for genome-wide barriers to persist at migration levels that would swamp individual adaptive alleles. An interesting question is how thresholds for congealing depend on the history of diverging populations, and whether certain population histories (e.g., involving rapid polygenic adaptation from standing genetic variation, as considered here) may lead to the emergence of genomewide congealing and RI over much shorter timescales than in the mutation-limited case (see also [47] for a discussion of rapid evolution of RI).

For local adaptation to be possible despite high migration (i.e., beyond the single-locus critical migration threshold) in our model, allele frequency divergence per locus must be significant (i.e., *p*_*c*_ − *p*_*r*_ *>* Δ_*c*_): this is a consequence of the rather extreme form of divergent selection, involving multiplicative fitness costs across loci, that we consider. In a more realistic setting with stabilizing selection towards different optima across different habitats, substantial divergence at the level of the quantitative trait may evolve even with very little differentiation at the underlying trait loci [25, 26, 48]. It is less clear whether the positive feedback between increasing trait divergence and falling effective migration rates that is so prominent under multiplicative selection also plays a role in this case, and how this depends on the genetic architecture of traits. Extending the approximations developed here to the case of stabilizing selection thus remains an interesting direction for future work.

### Effect of genetic drift on critical migration thresholds

Critical migration thresholds may be significantly reduced by drift when the loci underlying local adaptation have modest (*Ns*10) effects (see, e.g., Fig 1B, 2C, and 2D; also [23]), and when one habitat is much rarer than the other (*ρ* small). Moreover, the magnitude of this reduction is not very sensitive to *Ls*, with drift depressing critical migration thresholds by roughly similar amounts in both the LD-independent and the LD-dominated parameter regimes (i.e., for both the constant as well as the rising parts of the curves in Figs 2C and 2D).

Since direct estimation of the effect sizes of loci underlying adaptive divergence is difficult, one might turn the argument around and ask: given typical estimates of migration– between 1 and 10 ‘effective’ number of migrants exchanged per generation (see e.g., [49]), how strongly selected would a locus need to be to sustain adaptive divergence in the face of gene flow? Assuming *ρ*=0.1 in our model, adaptive divergence at a single locus requires *Ns>*9.7 with *Nm*=10 and *Ns>*2.05 with *Nm*=1. One might also look at an example where there is a modest genomewide barrier effect— with the same habitat proportions (*ρ*=0.1), *L*=100 and *s*=0.01 (so that *Ls*=1), we require *Ns>*7.6 with *Nm*=10 and *Ns>*1.7 with *Nm*=1 for adaptive divergence. The fact that the threshold *Ns* only decreases by a factor of 5 when *Nm* decreases by a factor of 10 (so that the threshold *m/s* is lower at lower *Nm*) is a consequence of genetic drift.

More broadly, this suggests that even though the effect sizes of loci contributing to local adaptation will depend on the size *N* of local demes, the *shape* of the distribution of effect sizes depends only on the rate of local drift relative to migration, i.e., on *Nm*. Thus, if smaller demes are connected by higher levels of migration, so that *Nm* is roughly similar across populations with very different local deme sizes, then critical *m/s* thresholds and the shape of the effect size distribution (on the *Ns* scale) would also be very similar across populations. Conversely, if smaller demes are also more cutoff from each other on average (i.e., have lower *Nm*), then the distribution of adaptive differences should be biased towards larger effect loci (on the *Ns* scale) in populations with smaller local demes.

### Effect of adaptive divergence on genome-wide neutral *F*_*ST*_

A striking observation that emerges from the analysis of the infinite-island model is that neutral, genomewide *F*_*ST*_ is actually highest for a pair of subpopulations belonging to the rare habitat (Fig 3B), even though there is *no* RI (and no isolation-by-distance) between them. This counter-intuitive finding can be rationalised by noting that *F*_*ST*_ between a pair of demes (within a larger mosaic of interconnected demes) does not measure the extent of *direct* genetic exchange between them, but instead depends on the level of exchange between an individual deme and the whole population (averaged over the two demes). Since demes in the rare habitat are more effectively isolated from the larger population than those in the common habitat, it follows that the average level of genetic exchange between any two randomly chosen demes and the population as a whole is lowest if both demes are in the rare habitat, and highest if both are in the common habitat, so that 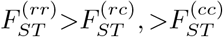 when habitats are at least partially locally adapted.

The fact that neutral diversity in the rare habitat is reduced much more strongly than in the common habitat is, to some extent, a consequence of the infinite-island setting, in which individuals are assumed to migrate from any island to any other with equal probability, irrespective of their habitat of origin or destination. One could consider alternative models, where islands are embedded in a 2D spatial matrix, and where habitats are spatially sorted, so that migration between islands belonging to the same habitat is more likely than between islands belonging to different habitats (models with habitat choice but no explicit space might result in qualitatively similar outcomes). However, as long as spatial sorting of habitats and/or habitat choice is not complete, we expect effective immigration into the rare habitat to be at least somewhat lower than into the common habitat, and average *F*_*ST*_ between demes belonging to the rare habitat to be accordingly highest, despite the lack of RI between them.

In their re-analysis of so-called genomic islands of divergence across various hybridizing populations, Cruikshank and Hahn (2014) [50] pointed out that increased *F*_*ST*_ may reflect reduced diversity (e.g., due to purifying selection within populations) rather than increased divergence (due to reduced gene flow between populations). A consideration of local adaptation in extended populations yields yet another caveat: where sampled subpopulations are embedded within a larger metapopulation, reduced gene flow between subpopulations belonging to different ecological niches can manifest itself primarily via reduced diversity within, rather than increased divergence between, subpopulations (inset, Fig 3B). Moreover, this reduction is more severe for subpopulations which support the niche that is more marginal or less abundant in the population as a whole. While this effect would be less extreme in reality than it is under the infinite-island model, it nevertheless points to the perils of neglecting the wider spatial context of two (or a few) sampled subpopulations when interpreting relative divergence between them. Note that this caveat also applies to markers linked to barrier loci (since the underlying argument is based only on there being a rare and a common habitat), and thus is also relevant to the interpretation of solitary peaks of *F*_*ST*_.

### Approximating the effects of LD via effective migration rates

The theoretical approximations developed here rely on two kinds of separation of timescales. First, we assume that immigrant genomes split up (via recombination) over timescales that are much shorter than those associated with the evolutionary dynamics of individual loci. Then LD *within* any subpopulation is weak, even when there is substantial LD at the level of the metapopulation as a whole. Thus, we can use single-locus (diffusion) results for allele frequencies, while approximating the effects of metapopulation-wide LD through an effective migration rate. In general, we expect this assumption to be valid if loci are unlinked or possibly weakly linked (and as long as *s, m*, 1*/N* « 1). By contrast, if deleterious alleles are tightly linked over a block of genome, then the effective unit of selection is the full block (rather than individual loci), and allele frequencies are shaped essentially by the balance between *m* and *Ls* [5].

Another factor that could affect the separation between timescales associated with single-locus and multi-locus dynamics is epistasis. For a given *F*_1_ fitness, diminishing returns epistasis between deleterious alleles (wherein selection per deleterious allele is weaker when many such alleles act in combination) tends to further strengthen the barrier to gene flow. Conversely, synergistic epistasis (alleles more deleterious in combination), which emerges naturally in models of stabilizing selection on additive traits, weakens the barrier relative to multiplicative selection [13]. While the prevalence of diminishing-returns epistasis in nature is unclear [51], it represents an interesting scenario (associated with potentially strong barrier effects) where our approximations may break down due to significant negative LD within subpopulations. Thus, generalizing the present analysis to arbitrary patterns of epistasis, including various forms of cryptic epistasis (which can generate strong RI even in the absence of strong ecological differentiation; see [52]), remains an interesting direction for future work.

The second key assumption is that even when drift has appreciable effects on allele frequencies, it has no effect on the dynamics of genotypes with multiple selected alleles, allowing us to approximate the latter via a *deterministic* effective migration rate. This second kind of *separation of timescales* emerges naturally when local adaptation is polygenic: selection per locus must then be at least as strong as drift for local adaptation to be possible, i.e., 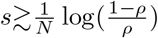, when loci are unlinked [38]. However, drift must then be much weaker than net selection against introgressing genotypes that carry multiple deleterious alleles, i.e., 1*/N* ∼ *s* « *Ls*. This allows us to use fundamentally different mathematical descriptions (deterministic vs. stochastic) for multi-locus and single-locus evolutionary dynamics. In practice, our approximations are accurate for even modestly polygenic architectures (e.g., see the *L*=10 plots in Figs 1A, 2A, and 2B).

The basic approach of approximating the effects of LD via an effective migration rate *m*_*e*_ is particularly useful for divergence based on quantitative traits, since *m*_*e*_ then depends primarily on the RV of migrants (see above). This can be estimated in the field, e.g., when pedigrees are available for ∼ 10 generations (see e.g., [34], [53]), thus providing (in principle) another estimate of the strength of the genomewide barrier to gene flow. The interpretation of effective migration rates in terms of RVs also suggests that at least for highly polygenic architectures of local adaptation, the gross effects of multi-locus LD on allele frequencies may depend on very few quantities, e.g., the mean fitness of *F*_1_ individuals and their within-family variance, even with an arbitrary effect size distribution [54]. Generalizing approximations based on effective migration rates to diploidy, unequal effect sizes and arbitrary dominance is a promising direction, as it will allow us to understand how hybridisation outcomes are influenced by selective interference between loci under different kinds of selective constraint, e.g., loci that are divergently selected (which may generate hybridisation load) and loci under background selection (which may contribute to heterosis).

## Acknowledgments

I thank Joachim Hermisson, Nick Barton and Sam Yeaman for very helpful comments on the manuscript.

## Supporting information

**S1 Text**. Supplementary Information

## SUPPLEMENTARY INFORMATION

### Section 1: Effective migration rates (neglecting drift)

#### Effective migration rate for deleterious alleles

We can solve eq. 3 of the main text iteratively to obtain *P*_*y*_, the deterministic frequencies of genotypes that carry *y*=1, 2, … *L* deleterious alleles, in a population at migration-selection equilibrium. These can be expressed as:

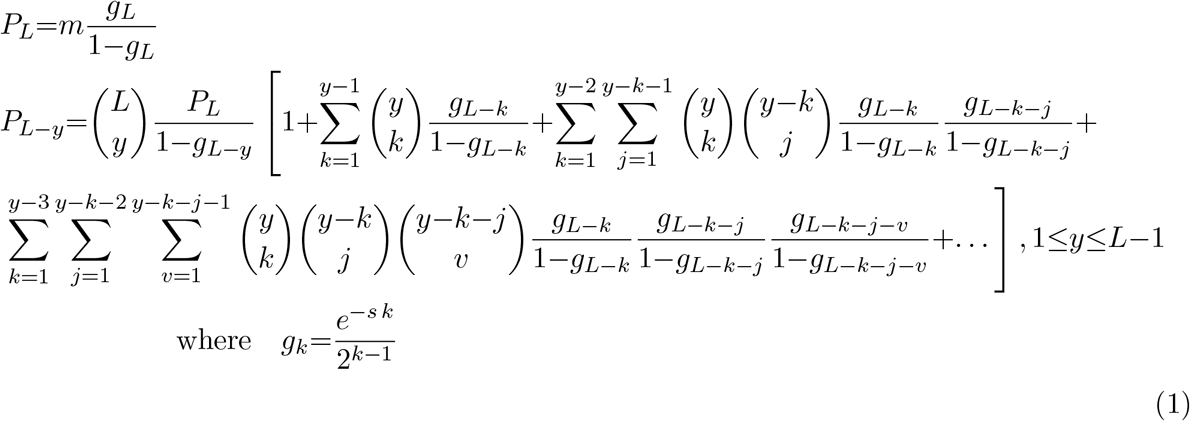

The above equation expresses the genotype frequency *P*_*L−y*_ for *y*=1, 2, … *L*−1 as a finite sum over *y* terms (i.e., the series within the square brackets contains 1, 2, 3 … terms in the expressions for *P*_*L−*1_, *P*_*L−*2_, *P*_*L−*3_, … respectively). It is also useful to represent these as infinite sums by Taylor expanding the various *g*_*k*_ */*(1−*g*_*k*_) in powers of *g*_*k*_. Note that 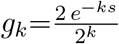 has a simple interpretation: it is the average number of offspring of an individual with *k* deleterious alleles, who inherit none of the *k* alleles; alternatively, it is the average number of offspring who inherit all *k* alleles. Then, we have the alternative representation:

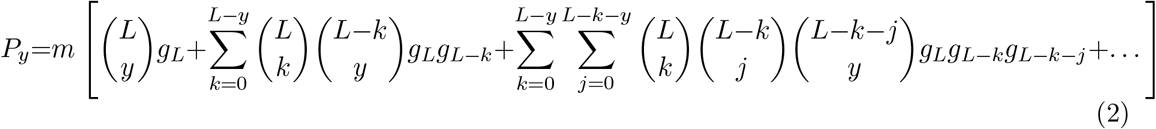

Note that the above representation involves an *infinite* series; the *k*^*th*^ term in the series is a *k*-fold convolution. The average deleterious frequency 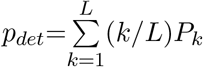 can be obtained by using either eq. (1) or (2). Here, we use eq. (2), thus expressing *p*_*det*_ as an infinite sum:

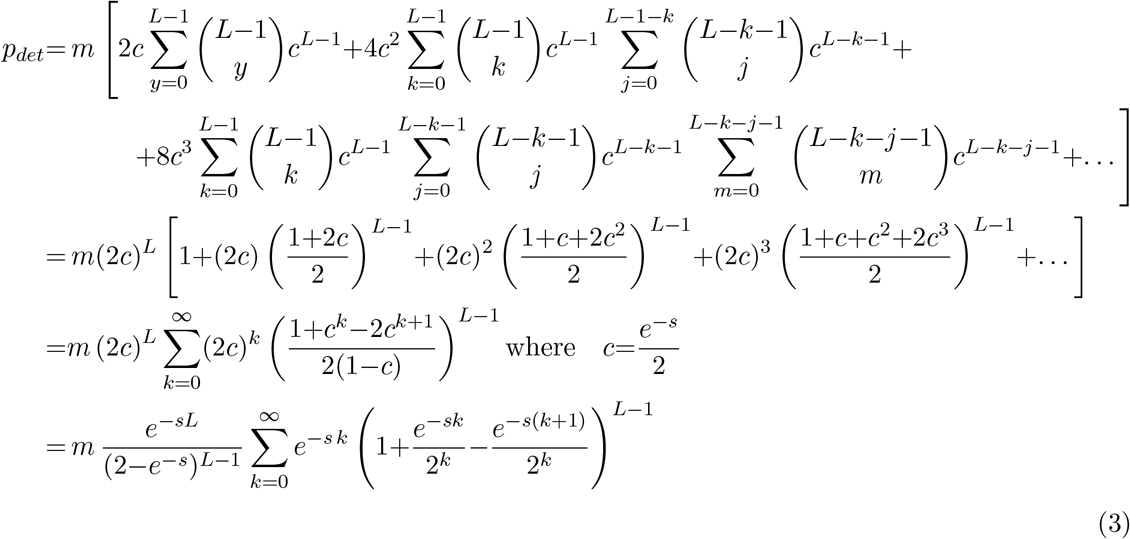

For *L*=1, the above expression reduces to: *p*_*det*_(*L*=1)=*m/*(*e*^*s*^−1) ≈ *m/s*. Following the main text, we define the effective migration rate *m*_*e*_(*s, L*) as the migration rate which would cause the deleterious allele frequency at a single locus under migration-selection equilibrium (with selection coefficient *s* for the deleterious allele) to be equal to *p*_*det*_, which is the average equilibrium frequency that emerges in the multi-locus model, in which immigrant genotypes carrying *L* such deleterious alleles are introduced at rate *m* per generation. Then, we have: *m*_*e*_(*s, L*)=*p*_*det*_(*e*^*s*^−1)≈*sp*_*det*_, where *p*_*det*_ is given by eq. (3). This finally yields the expression for *m*_*e*_(*s, L*) in eq. 4 in the main text.

In the limit *s*→0, *L*→∞ with *s*(*L*−1)=*θ* held constant, we can approximate *m*_*e*_(*s, L*)*/m* as:

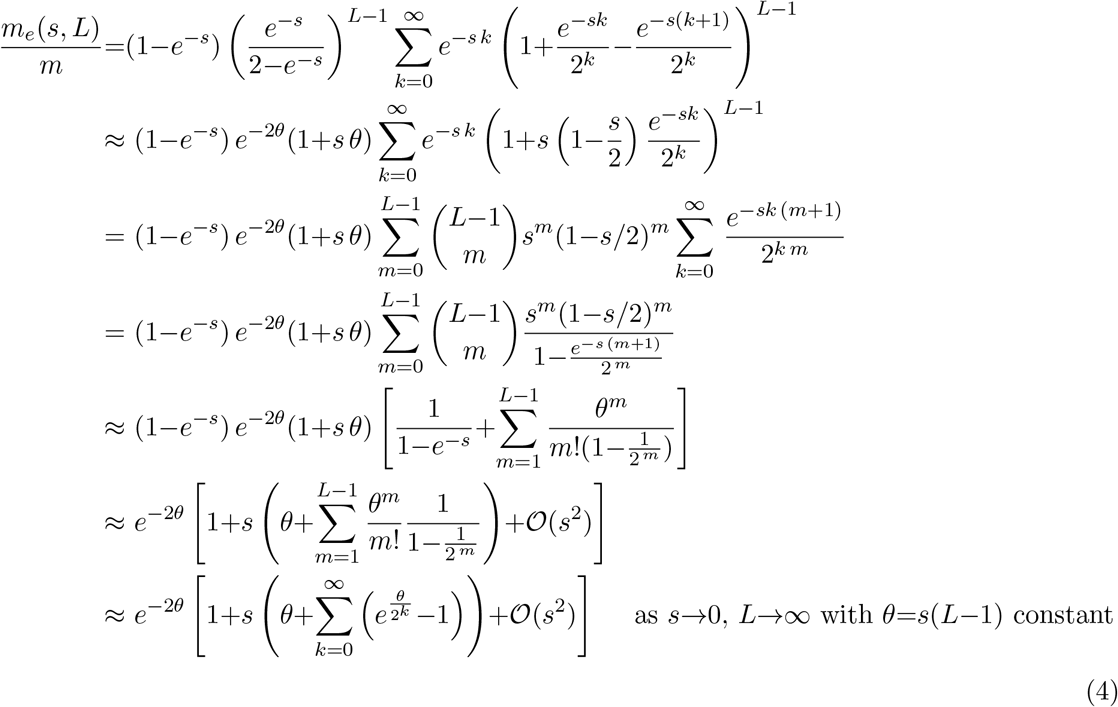

This is approximated reasonably well by: 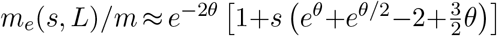. Thus, the effective migration rate of selected alleles is reduced relative to the raw migration rate by a factor that converges (in the highly polygenic limit) to the average RV *e*^*−*2*sL*^ of migrants (see also the explanation following eq. 5 of the main text).

### Effective migration rate for neutral alleles

Following Bengtsson (1985), we define the effective migration rate 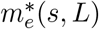 associated with a neutral locus to be the rate at which neutral alleles embedded within incoming migrant genomes (which carry *L* unlinked deleterious alleles of effect *s*) are transferred via one or more recombination events onto the wildtype background (which carry no deleterious alleles) in a population at migration-selection equilibrium.

To calculate this, we must first calculate the equilibrium frequencies {*f*_*y*_} of genotypes that carry the neutral allele in conjunction with *y* deleterious alleles (each with selective effect *s*), where all alleles (neutral and deleterious) are unlinked. As before, we assume that the frequency of any genotype with deleterious alleles is sufficiently low that quadratic terms of the kind *f*_*y*_*f*_*y*_*′* can be neglected. Then *f*_*y*_ satisfy the following equations in steady state:

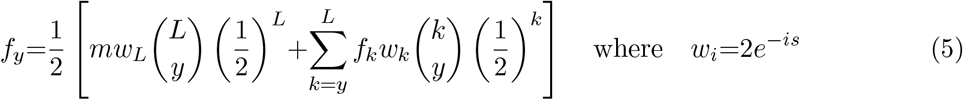

Note that these are very similar to eq. 3 in the main text, except that the right hand side now has an additional factor of 1*/*2, which is the probability that an individual carrying the neutral allele transmits it to an offspring. Equation (5) can be solved as before; it is again useful to express the various *f*_*y*_ as an infinite series in powers of *g*_*k*_, so that we have:

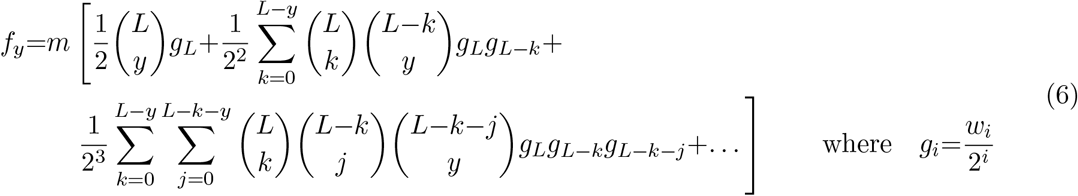

The above expression allows us to read off the distribution of *t*_*esc*_, the time taken for the focal (neutral) allele to ‘escape’ the deleterious background, by setting *y*=0. Thus, we have:

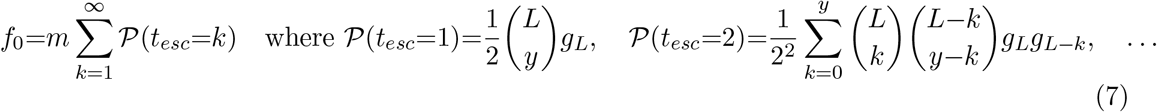

These can be used to calculate various moments 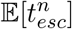 of the time to escape.

We can also use {*f*_*y*_} to obtain the effective migration rate 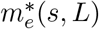 for neutral alleles using:

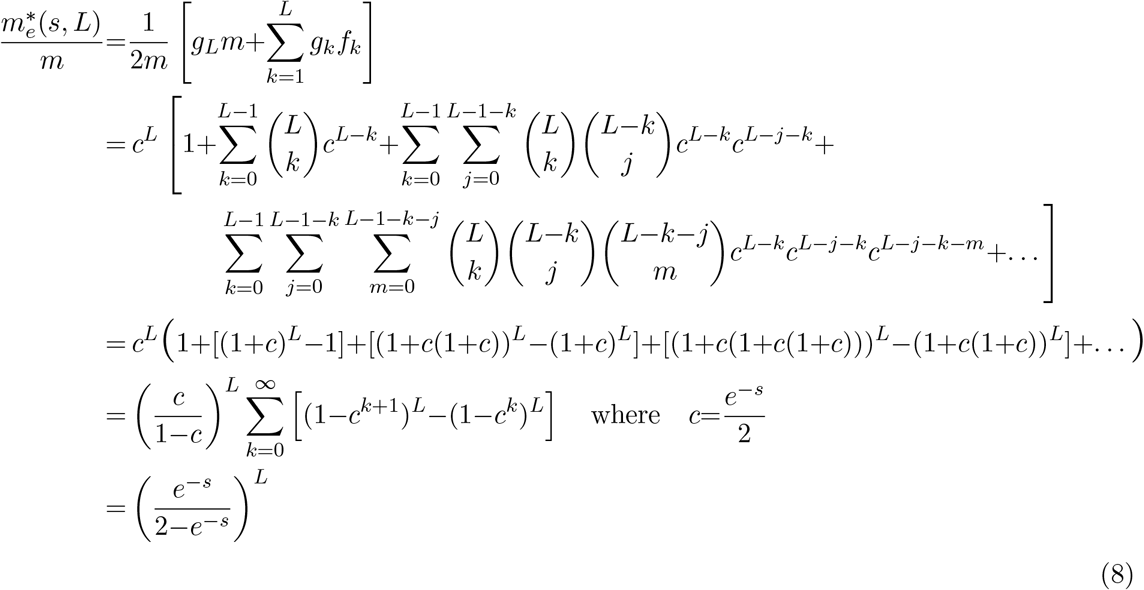

In the highly polygenic limit *s*→0, *L*→∞ with *sL*=*θ*_*∗*_ held constant, this simplifies to 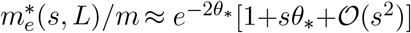.

### Section 2: Calculating average coalescence times and expected *F*_*ST*_ in the infinite-island model using the structured coalescent

To calculate coalescence times, it is useful to consider a population with a finite number *D* of demes, and then take the *D*→∞ limit. There are 5 distinct possibilities for randomly sampling 2 lineages from *D* demes, where a fraction *ρ<*1*/*2 of demes belong to the rare habitat and a fraction 1−*ρ* to the common habitat. With probability *ρ/D*, both lineages will be sampled from the *same* deme (arbitrarily labeled *i*) in the rare habitat; with probability (1−*ρ*)*/D*, both will be from the *same* deme *i* in the common habitat; with probability *ρ*(1−1*/D*), lineages will be sampled from two *different* demes (arbitrarily labeled *i* and *j*) both in the rare habitat; with probability (1−*ρ*)(1−1*/D*), lineages will be from two different demes *i* and *j* both belonging to the common habitat; with probability 2*ρ*(1−*ρ*), lineages will be sampled from two different demes *i* and *j*, one belonging to the rare and the other to the common habitat. We denote the 5 coalescence times corresponding to these 5 different configurations for 2 lineages by: *T*_*i,i*; *r*_, *T*_*i,i*; *c*_, *T*_*i,j*; *r,r*_, *T*_*i,j*; *c,c*_, and *T*_*i,j*; *r,c*_.

The expected values of the various *F*_*ST*_ measures (introduced in the main text) can be expressed in terms of the expected coalescence times as follows:

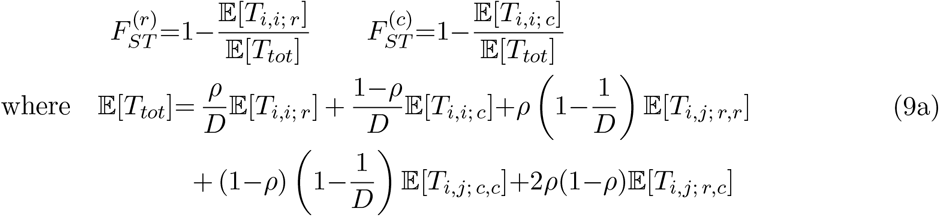

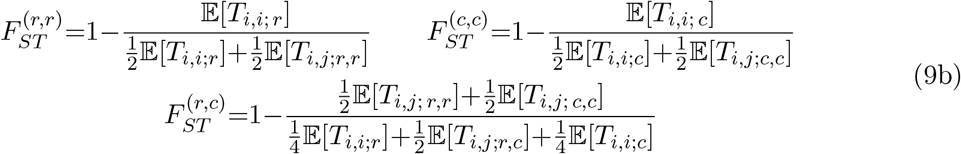

In order to calculate the expected coalescence times, we consider how lineages trace back in time through different demes belonging to one or other habitat. Let *m*_*rc*_ and *m*_*rr*_ denote the probability per unit time that a lineage in a deme within the rare habitat traces back to the rare and common habitats respectively. We can define analogous backward migration rates *m*_*cc*_ and *m*_*cr*_ for lineages in demes belonging to the common habitat. We use uppercase letters to denote the corresponding population-size-scaled migration rates *M*_*rr*_=*Nm*_*rr*_, *M*_*rc*_=*Nm*_*rc*_, *M*_*cc*_=*Nm*_*cc*_, *M*_*cr*_=*Nm*_*cr*_. Note that the rates *m*_*rr*_ and *m*_*cc*_ (or *M*_*rr*_ and *M*_*cc*_), as defined here, include backward migration events where a lineage emigrates from a deme but then immigrates back into the same deme (though this occurs with a small probability 1*/D*, which vanishes in the limit *D*→∞).

Going back one generation into the past, there are three possibilities for a pair of lineages that are, at present, in the same deme within the rare habitat: the two lineages can coalesce with probability 1*/N* ; one or other of the two lineages can trace back to a different deme within the rare habitat with probability 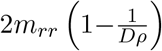 (here, the factor 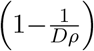 accounts for the fact that the migration rate *m*_*rr*_ also includes ‘migration’ events which lead a lineage back to its deme of origin); finally, one or other lineage can trace back to a deme belonging to the common habitat with probability 2*m*_*rc*_. Note that we neglect events involving simultaneous migration of both lineages in a single timestep as these occur with vanishingly small rates under the usual coalescent scaling. Similarly, by enumerating all possible single-generation events for the other 4 two-lineage configurations, we arrive at the following set of recursions for the expected coalescence times:

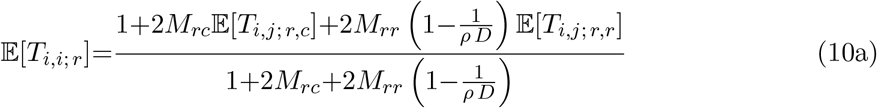

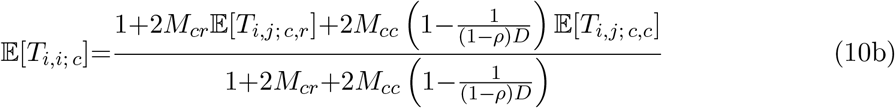

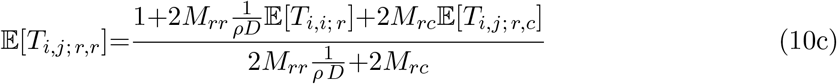

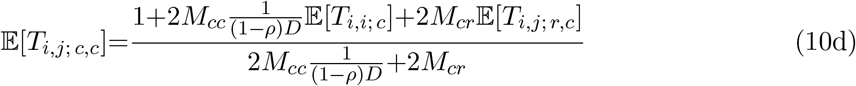

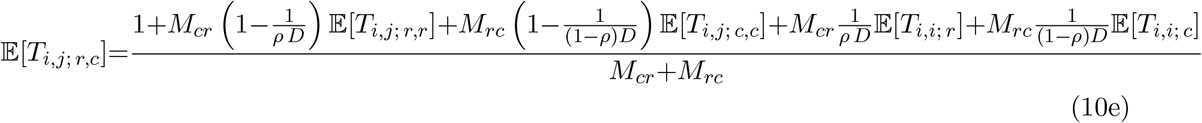

Solving equation 10 for the expectations of the 5 coalescence times, substituting these expressions into eq. (9), and then taking the limit *D*→∞ (which allows us to neglect all terms that are 𝒪 (1*/D*)) finally gives the expected values of the various *F*_*ST*_ measures (eq. 9 of the main text).

### Section 3: Convergence to the small *s* (large *L*) limit

As shown in Section 1, in the highly polygenic limit *s*→0, *L*→∞ with *θ*=(*L*−1)*s* or *θ*_∗_=*Ls* fixed, the (scaled) effective migration rates *m*_*e*_(*s, L*) and 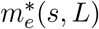 converge to *m*_*e,∞*_(*s, L*) = *m e*^*−*2*θ*^ and 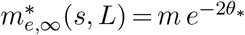 respectively. Note that in this limit, we also have *θ*→*θ*_∗_. However, I still distinguish between the two to highlight the conceptual distinction that the barrier effect at any selected locus is due to the other *L*−1 selected loci, while the barrier effect at a neutral locus is due to *L* selected loci.

As discussed in Section 1, for fixed *θ* or *θ*_∗_, the lowest order (in *s*) correction to the asymptotic prediction gives (see also eq. 6 of the main text):

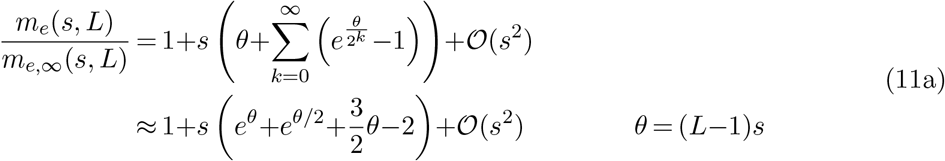

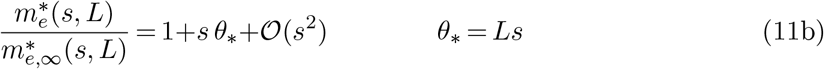

Figure 1 illustrates this convergence by plotting (*m*_*e*_*/m*_*e,∞*_)−1 for selected alleles (filled symbols) and 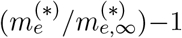 for neutral alleles (empty symbols) vs. *L* for different values of *θ* or *θ*_∗_ (different colors). The small *s* (or equivalently, large *L*) approximations in equations (11a) and (11b) are shown using solid and dashed lines. As expected, both effective migration rates converge towards the highly polygenic prediction as *L* increases (or equivalently, *s* decreases). Moreover, this convergence is captured quite accurately by the approximations above.

Note that for a given *Ls*, the barrier effect is weaker, i.e, *m*_*e*_ higher, when the total selective disadvantage is due to a small number of loci of relatively strong effect. Conversely, the barrier effect is strongest when the total selective disadvantage is due to very many infinitesimal loci.

One can also ask: to what extent is local adaptation sensitive to *L* (or *s*) for a given total selective disadvantage *Ls*, i.e., (when) does the detailed genetic architecture of local adaptation matter? Figure 1b shows theoretical predictions for the frequency of the locally adaptive allele on the island vs. *m/s* for various numbers *L* of divergently selected loci (various colors), for fixed *θ*=(*L*−1)*s*, and fixed *Ns*. The finite *L* predictions are obtained by solving eq. 7 of the main text, using the expression for *m*_*e*_ in eq. 4, while the *L*→∞ predictions use the asymptotic expression *m*_*e*_(*s, L*)=*m e*^*−*2*θ*^ instead of eq. 4. Figure 1b shows these plots for *θ*=0.5 (solid lines), *θ*=1.0 (dashed lines) and *θ*=2 (dotted lines). We see that allele frequencies are more sensitive to the actual genetic architecture (i.e., to *L* and *s* individually) when *θ* is large (dotted lines). Conversely, for smaller *θ*, local adaptation depends on *s* only via the composite parameters *Ls, Ns* and *m/s*.

**Figure 1:**
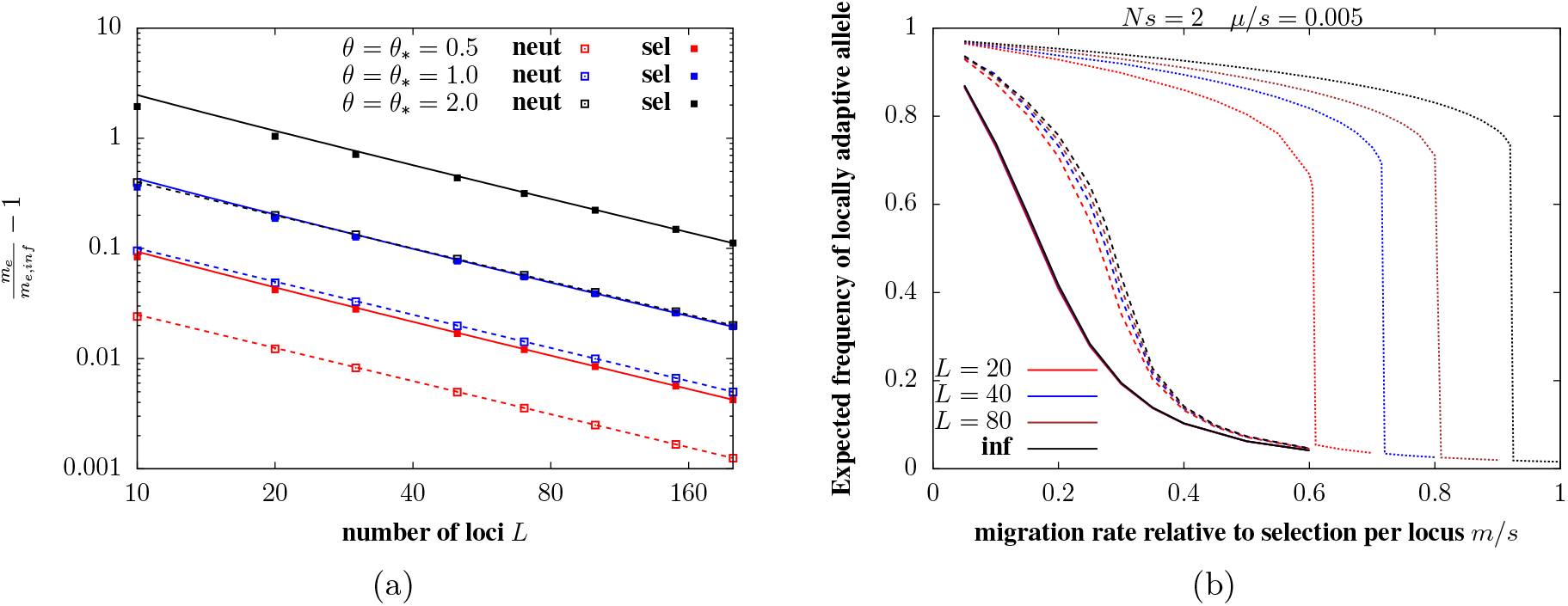
(a) Relative deviations *m*_*e*_*/*(*m*_*e,∞*_)*−*1 (filled squares) and 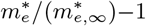 (empty squares) vs. *L*, the number of selected loci, where *m*_*e*_ and 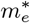 are the effective migration rates associated with deleterious and neutral alleles (calculated using eqs. 4 and 5 of the main text), and *m*_*e,∞*_ = *m e*^*−*2*θ*^ and 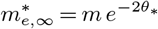 the corresponding rates in the highly polygenic limit, obtained by taking *s→*0, *L→∞*, with *θ*=*s*(*L−*1) and *θ*_∗_=*sL* held constant. The different colors correspond to different values of *θ* or *θ*_∗_. Solid and dashed lines show approximate expressions for these deviations (equations (11a) and (11b)), which are correct to first order in *s*. (b) The expected frequency of the locally adaptive allele on the island vs. *m/s*, under mainland-island migration. Solid, dashed and dotted lines correspond to *θ*=0.5, 1 and 2 respectively; the different colors correspond to different values of *L*; other parameters are *Ns*=2 and *µ/s*=0.001. All finite *L* predictions in (b) are obtained by numerically solving eq. 7 in conjunction with eq. 4 of the main text; the predictions for highly polygenic architectures (in black) are obtained by numerically solving eq. 7 while using *m*_*e,∞*_=*e*^*−*2*θ*^ for the effective migration rate.

In fig. 1b, *s* is decreased (while also simultaneously decreasing *m* and *µ* and increasing *L* and *N*) in order to keep the scaled parameters *Ns, m/s, µ/s* and the total selective effect *Ls* constant, since the goal is to illustrate how in the highly polygenic limit, evolutionary outcomes depend on *s* only via these composite parameters. However, in order to investigate how evolutionary outcomes depend on the genetic architecture of local adaptation for a given total selective disadvantage *Ls* in a population of a certain size *N* and subject to a certain rate of migration *m*, we must compare scenarios where lower *s* implies lower *Ns* and higher *m/s*. Thus, even though more polygenic architectures can, in principle, generate stronger multi-locus barrier effects (see, e.g., fig. 1a), they are also associated with reduced efficacy of selection and increased swamping at individual loci (via reduced *Ns* and increased *m/s*), making local adaptation (in our model) more difficult for more polygenic architectures.

### Section 4: Effect of mutation rate on local adaptation in the mainland-island model

In the main paper, the mutation rate for the mainland-island model was held fixed at *µ/s*=0.005. Figure 2 contrasts adaptation thresholds shown in the main paper with those for *µ/s*=0.00125 (mutation rate lower by a factor of 4), for two different population sizes corresponding to *Ns*=2 and *Ns*=8. We see that theoretical predictions that account for both LD and drift (obtained from eq. 7 in conjunction with eq. 4 of the main paper) are quite accurate, also for the lower mutation rate (lines vs. symbols in fig. 2)

**Figure 2:**
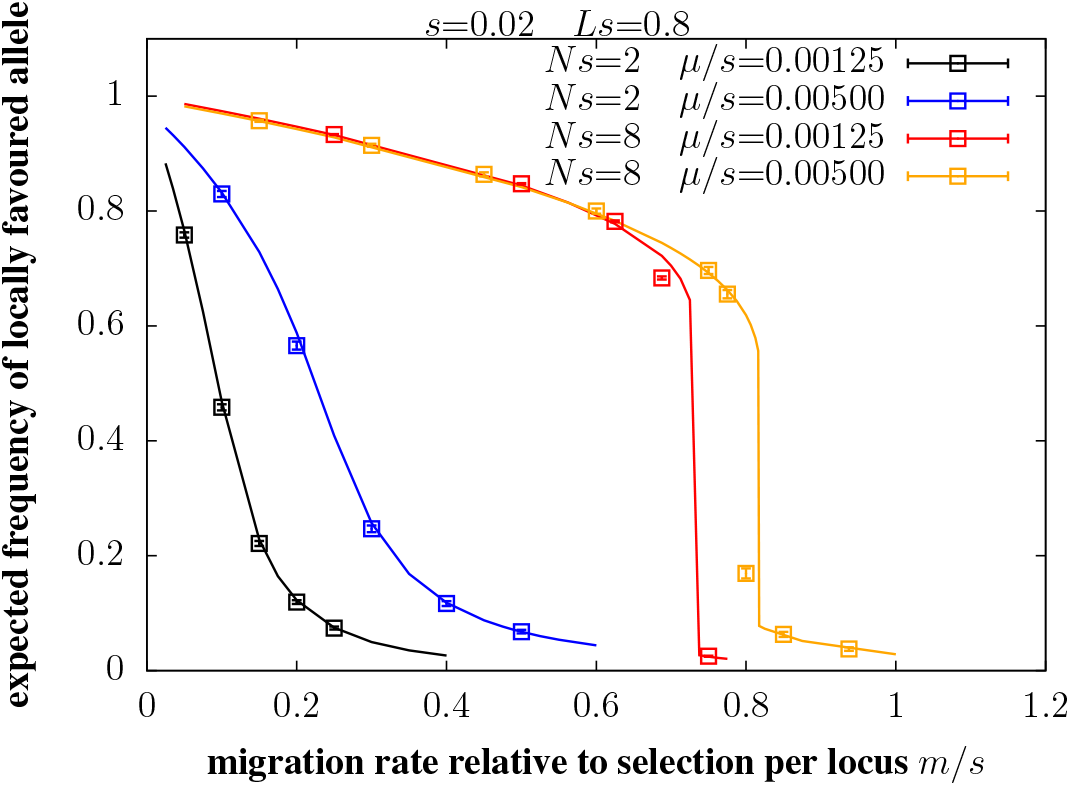
Expected frequency of the locally adaptive allele on the island vs. *m/s*, migration rate relative to selection per locus for two different population sizes (*Ns*=2 and *Ns*=8) and two different mutation rates (*µ/s*= 0.00125 and 0.005) for each. Other parameters are: *s*=0.02, *Ls*=0.8. Results of individual-based simulations (symbols) are in good agreement with theoretical predictions (lines) obtained from eqs. 7 and 4 of the main text for both mutation rates.

As expected, local adaptation is more sensitive to mutation in the smaller (*Ns*=2) population, with adaptive allele frequencies being significantly lower for the lower mutation rate across all migration levels. Thus, in the smaller population, mutation is instrumental in countering the loss of local adaptation due to drift and gene flow. By contrast, in the larger (*Ns*=8) population, the adaptive allele frequency is actually marginally higher for the lower mutation rate for *m/s*; ≲0.5. Thus, in this regime, mutation has very little effect on polymorphism unless mutation rates are extremely small; e.g., *µ/s* must be at least as low as ∼ 10^*−*6^ for *m/s*=0.2, and ∼ 2×10^*−*5^ for *m/s*=0.4, to have an appreciable effect on the adaptive allele frequency in the larger population. Sensitivity to mutation rates does, however, increase closer to the threshold for loss of local adaptation, with the threshold *m/s* falling by about 12% when the mutation rate is reduced from 0.005 to 0.00125. We also find that local adaptation is insensitive to mutation rates over a wider range of *m* when *Ls* is larger, as effective migration rates (which govern the extent of maladaptive gene flow) can then be much lower than *m*.

### Section 5: Deterministic analysis of local adaptation in the infinite-island model

As in the main paper, let *p* denote the frequency of the allele that is advantageous in the common (and disadvantageous in the rare) habitat. In the deterministic limit, i.e., neglecting drift, we can write down coupled equations for the time evolution of *p*_*r*_ and *p*_*c*_, the allele frequencies in the rare and common habitats:

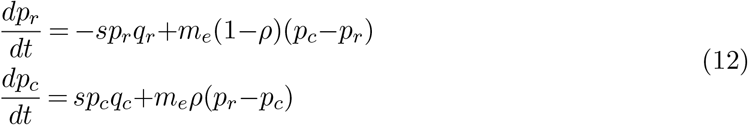

The assumption here is that allele frequency evolution at any locus is governed by the balance between selection at that locus and migration which is described by an effective migration rate *m*_*e*_, which captures the ‘barrier effect’ due to selection against deleterious alleles at the other *L*−1 loci. We will use Δ=*p*_*c*_−*p*_*r*_ to denote the allele frequency difference between the two habitats, and express the effective migration rate as *m*_*e*_= *m g*[Δ]. Here, *g*[Δ], which is the gene flow factor, is assumed to depend on the allele frequencies *p*_*c*_ and *p*_*r*_ only via the difference Δ (see also main paper).

One can employ various approximations for *g*[Δ] (see below) and solve eq. (12) numerically to obtain *p*_*c*_ and *p*_*r*_. The deterministic predictions for *m*_*c*_*/s* (the threshold for local adaptation in the rare habitat) in figs. 2c and 2d in the main text are obtained in this way— by using 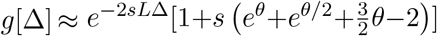 (see eq. (11a)), and then numerically solving (12) at equilibrium (i.e., with *dp*_*c*_*/dt*=*dp*_*r*_*/dt*=0), to determine the value of *m*_*c*_*/s* beyond which no non-zero solution for Δ exists. Where multiple equilibria are possible, one can use eq. (12) to determine which equilibrium is reached by numerically solving for *p*_*r*_(*t*) and *p*_*c*_(*t*) as a function of time *t*, given initial allele frequencies *p*_*r*_(0) and *p*_*c*_(0).

We can also obtain approximate analytical expressions for *m*_*c*_*/s* by assuming that the common habitat is always reasonably well-adapted, such that *q*_*c*_=1−*p*_*r*_ is small and 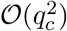 terms can be neglected. This allows us to approximate: *p*_*c*_*q*_*c*_≈*q*_*c*_ and *p*_*r*_*q*_*r*_≈Δ(1−Δ)+*q*_*c*_(1−2Δ), which gives the following coupled equations for the time evolution of *p*_*c*_ and Δ.

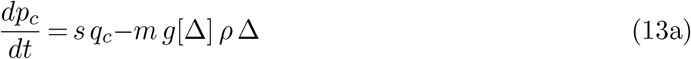

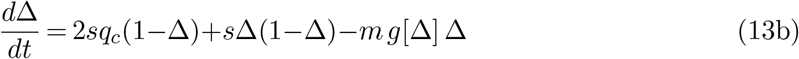

At migration-selection equilibrium (i.e., assuming the time derivatives in the above equation to be zero), we can use eq. (13a) to express *q*_*c*_ in terms of Δ, and then substitute into (13b) to obtain the following equation for Δ *at equilibrium*:

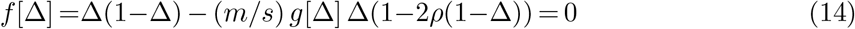

There is always an equilibrium at Δ=0 (corresponding to zero adaptive differentiation between habitats): this is unstable as long as *m/s<*(1−2*ρ*) and is stable otherwise. Note that *m*_*c*,1_*/s*=(1−2*ρ*) is simply the threshold for the maintenance of polymorphism at a single locus (or equivalently, the LE threshold) under deterministic migration-selection equilibrium.

In addition, there may be other equilibria at Δ=Δ_∗_*>*0 (corresponding to partial local adaptation in the rare habitat), where Δ_∗_ satisfies 1−Δ_∗_=(*m/s*)[1−2*ρ*(1−Δ_∗_)]*g*[Δ_∗_]. While this does not allow us to solve for Δ_∗_ as a function of *m/s*, we can obtain an explicit expression for the critical migration threshold (denoted by *m*_*c*,2_*/s*) beyond which no such equilibrium (with Δ_∗_*>*0) exists. As shown below, the threshold *m*_*c*,2_*/s* may or may not be equal to the LE threshold *m*_*c*,1_*/s*, depending (primarily) on the values of *Ls* and *ρ*.

As a prelude to deriving an expression for *m*_*c*,2_*/s*, it is useful to visualize the behaviour of *f* [Δ] as a function of Δ (see eq. (14)). For the purpose of this visualisation, the gene flow factor *g*[Δ] is approximated by its asymptotic large *L* form: *g*[Δ]≈*e*^*−*2*Ls* Δ^. However, the qualitative behaviours we observe are not sensitive to this assumption. Figures 3a and 3b show *f* [Δ] (scaled by *s*) vs. Δ for *Ls*=0.5 and *Ls*=1 respectively, for *ρ*=0.1, with the different colors in each plot corresponding to different values of *m/s*. Those points at which the curves intersect the horizontal axis (i.e., at which *f* [Δ]=0) correspond to equilibria. An equilibrium is *stable* if *f* [Δ] is a decreasing function of Δ, i.e., if the curve is downward sloping at the equilibrium, and is unstable otherwise.

For small *Ls* and low values of *m/s* (red and brown curves in fig. 3a), there are two equilibria— an unstable equilibrium at Δ=0 and a stable equilibrium at Δ=Δ_∗_*>*0 (marked by the × symbol on the plots). Increasing *m/s* causes Δ_∗_ to decrease; above a critical migration threshold *m*_*c*,1_*/s*=1*/*(1−2*ρ*), the equilibrium at Δ=0 becomes stable while the alternative equilibrium Δ_∗_ becomes unstable and also negative (and thus is no longer biologically meaningful). The transition threshold *m*_*c*,1_*/s*=1*/*(1−2*ρ*) (blue curve in fig. 3a), at which divergence between habitats collapses, corresponds to a so-called transcritical bifurcation.

**Figure 3:**
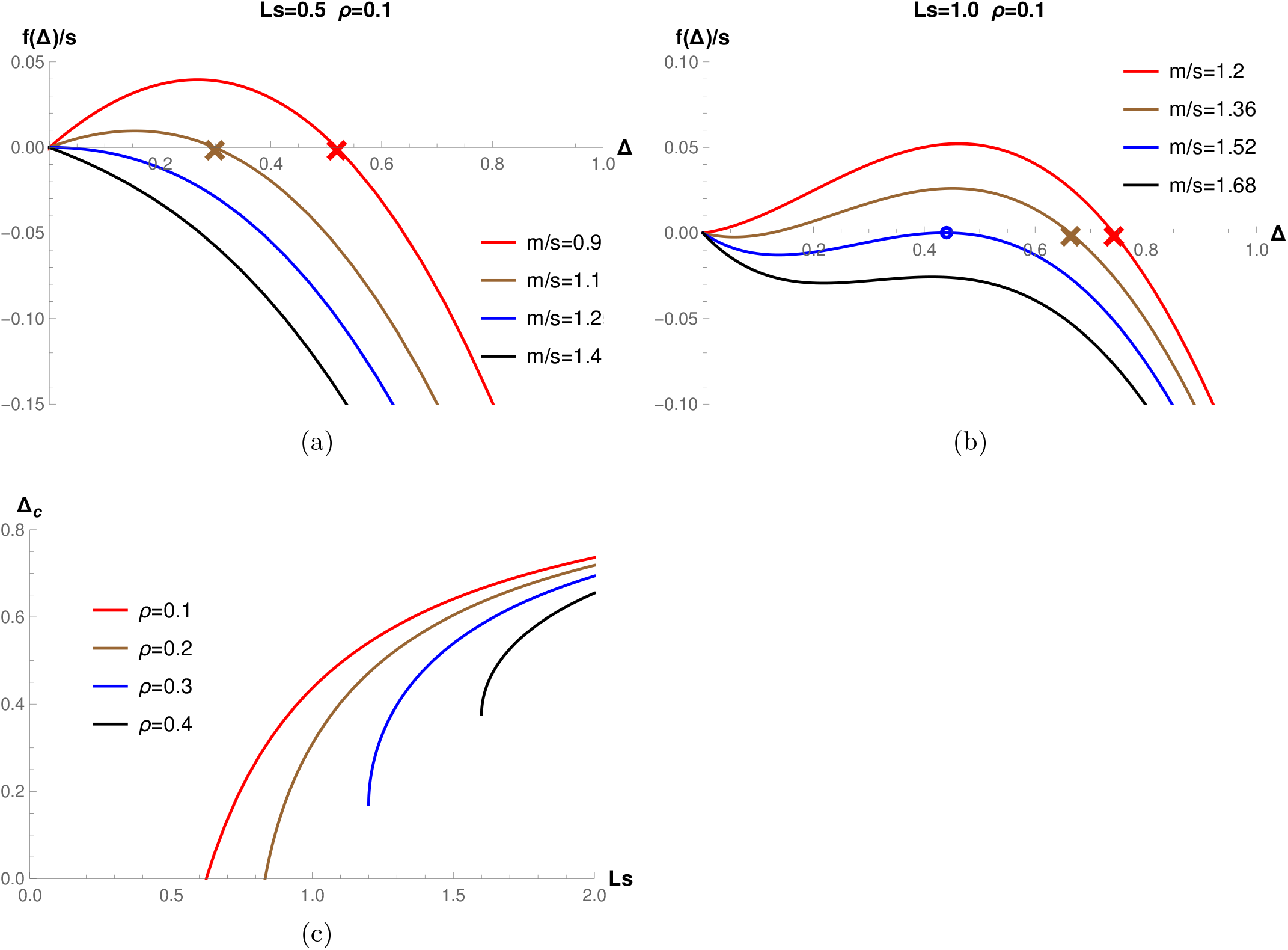
Deterministic analysis of local adaptation in the infinite-island model in the highly polygenic limit. (A)-(B) *f* [Δ] (scaled by *s*) vs. Δ for various values of *m/s* for *ρ*=0.1 and (A) *Ls*=0.5 (B) *Ls*=1. Here, Δ=*p*_*c*_*−p*_*r*_ is the allele frequency difference between the two habitats, and *f* [Δ], defined in eq. (13b), determines equilibria. Equilibria are those values of Δ at which *f* [Δ]=0; stable equilibria are those for which the curve *f* [Δ] vs. Δ is downward sloping, i.e., at which *df* [Δ]*/d*Δ is negative. For low *Ls* (fig. A), there is a critical migration level *m*_*c*,1_*/s*=1*/*(1*−*2*ρ*) (blue curve), above which the Δ=0 equilibrium becomes stable, while the Δ*>*0 equilibrium (marked by *×* symbols) vanishes. For high *Ls* (fig. B), there are two thresholds: the Δ=0 equilibrium becomes stable for *m/s>*1*/*(1*−*2*ρ*) but the Δ*>*0 equilibrium also persists, along with a third unstable equilibrium. At a second migration threshold *m*_*c*,2_*/s* (blue curve), the stable and unstable equilibrium collide: the corresponding value of Δ at which this occurs is Δ_*c*_ (marked by a circle in fig. B), which is the lowest possible divergence level that is stable (for *m>m*_*c*,1_). Above the migration threshold *m*_*c*,2_, only a single stable equilibrium at Δ=0 exists. (C) The critical adaptive divergence, Δ_*c*_ (obtained from eq. (16a)), vs. *Ls* for various *ρ*. There is a threshold (*Ls*)_∗_ (which increases with increasing *ρ*) such that Δ_*c*_ is non-zero only for *Ls>*(*Ls*)_∗_.

For large *Ls* (fig. 3b), we observe qualitatively different transitions of the equilibria with changing *m/s*. At low levels of migration, i.e., for *m/s <m*_*c*,1_*/s* = 1*/*(1−2*ρ*) (red curve), there is (as before) an unstable equilibrium at Δ=0 and a stable equilibrium at Δ=Δ_∗_*>*0. At intermediate migration levels (brown curve), i.e., for *m*_*c*,1_*/s <m/s <m*_*c*,2_*/s*, the Δ=0 equilibrium becomes stable; however, the stable Δ_∗_*>*0 equilibrium also persists, and there now appears a third (unstable) positive equilibrium separating the two stable equilibria. Increasing *m/s* now causes the unstable and stable equilibria to converge towards a common value Δ_*c*_. At a critical migration threshold *m*_*c*,2_*/s* (blue curve), the two equilibria collide (this corresponds to a so-called saddle-node bifurcation), and we have Δ_∗_=Δ_*c*_ (marked by a circle in fig. 3b). Above this threshold, the two non-zero (stable and unstable) equilibria vanish, and there exists only a single stable equilibrium at Δ=0 (black curve). Note that *m*_*c*,2_*/s* is greater than the threshold *m*_*c*,1_*/s* at which the Δ=0 equilibrium becomes stable.

We can determine Δ_*c*_ by noting that it is that value of Δ at which both *f* [Δ] and its first derivative with respect to Δ become zero (note the shape of the blue curve at this equilibrium). Using eq. (14), the two conditions— *f’
s*[Δ_*c*_]=0 and *f ’* [Δ_*c*_]=0— imply respectively:

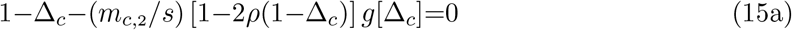

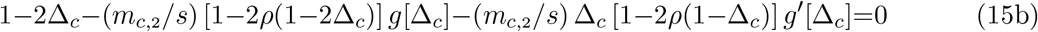

Consider first the highly polygenic limit (*s*→0, *L*→∞ and *Ls* constant), in which the gene flow factor can be approximated as *g’
s*[Δ] ≈ *e*^*−*2*Ls*Δ^, so that we have: *g*[Δ] = −2*Ls g*[Δ]. Substituting into eq. (15) yields a simple quadratic equation in Δ_*c*_, which involves only on *Ls* and *ρ*; solving this for Δ_*c*_ and then substituting into eq. (15a) gives *m*_*c*,2_*/s*. Thus, in this limit, we have the following expressions for the critical migration threshold and the critical divergence level:

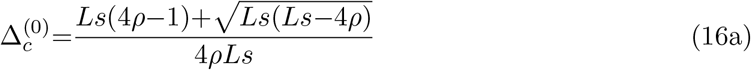

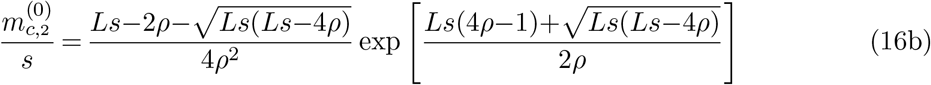

Figure 3c shows the prediction 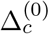 as a function of *Ls* for various values of *ρ*. Note that there exists a non-zero 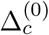 only above a threshold value of *Ls*, which we denote by (*Ls*)_*∗*_. For *ρ<*1*/*4, the critical divergence level 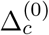 decreases as *Ls* decreases, approaching 0 as *Ls*→(*Ls*)_*∗*_. This allows us to solve for (*Ls*)_*∗*_ by setting 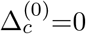 in eq. (16a), which gives (*Ls*)_*∗*_ =1*/*[2(1−2*ρ*)] for *ρ<*1*/*4. For *ρ>*1*/*4, the critical divergence level 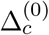 approaches a non-zero value as *Ls*→(*Ls*) _*∗*_: from eq. (16a), it follows that this bifurcation must occur at *Ls*=4*ρ*. Thus, for *ρ>*1*/*4, we have (*Ls*)_*∗*_ =4*ρ*, with 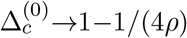 as *Ls*→(*Ls*)_*∗*_. This thus provides the deterministic prediction for the threshold (*Ls*)_*∗*_ above which LD can sustain local adaptation migration at rates higher than the single-locus critical migration threshold. This prediction for (*Ls*)_*∗*_ is shown using dashed vertical lines in figs. 2C and 2D of the main paper.

We can obtain a better approximation for *m*_*c*,2_*/s* and Δ_*c*_ (which is correct to first order in *s*) by using the expression in eq. (11a) for the gene flow factor. For generality, let us express this as: *g*[Δ]=*e*^*−*2*Ls*Δ^(1+*s g*_1_[Δ]); in all numerics, we will use *g*_1_[Δ]=*e*^*Ls*Δ^+*e*^*Ls*Δ*/*2^−2+ (3*/*2)Δ. Substituting this into equation (15) does not yield an exact explicit solution for Δ_*c*_ and *m*_*c*_*/s*. However, one can obtain an approximate solution using a perturbative expansion in *s*: this involves expressing *m*_*c*,2_*/s* and Δ _*c*_ as 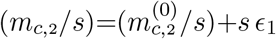 and 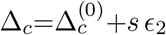, and then solving for the perturbation terms *ϵ*_1_ and *ϵ*_2_. This finally yields:

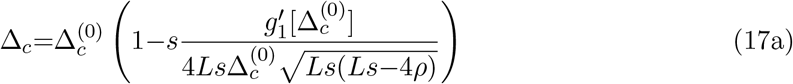

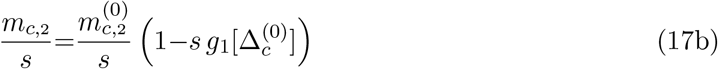

Figures 2C and 2D in the main paper show both the asymptotic (*L*→∞, *s*→0) prediction (eq. (16b)) for the critical migration threshold, and the more accurate (correct to first order in *s*) prediction in eq. (17b).

### Section 6: Dependence of critical migration thresholds on the initial state of the population

The existence of multiple stable equilibria (for *Ls>*(*Ls*)_*∗*_ and *m*_*c*,1_*<m<m*_*c*,2_) in the deterministic model above suggests that in this parameter regime, evolutionary outcomes in the stochastic model (i.e., with finite *Ns*) may also be quite sensitive to the *initial state* of the population, i.e., to allele frequency differences between demes and/or genetic variation within demes. We can gain insight into this by comparing simulations initialized in different ways, keeping all other parameters constant.

For simplicity, all loci in all demes in the rare habitat are initialized with the same allele frequency (denoted by *p*_*r*,0_); similarly, all loci in all demes in the common habitat are initialized with allele frequency *p*_*c*,0_. Figure 4 shows the results of individual-based simulations with 4 different initialization conditions (ICs): *p*_*r*,0_=0, *p*_*c*,0_=1 (IC1; red), *p*_*r*,0_=0.5, *p*_*c*,0_=0.5 (IC2; blue), *p*_*r*,0_=0.8, *p*_*c*,0_=0.8 (IC3; brown) and *p*_*r*,0_=0.9, *p*_*c*,0_=0.9 (IC4, orange). IC1 corresponds to a scenario of secondary contact where both habitats are perfectly adapted, resulting in maximum possible adaptive divergence and maximum possible barrier effects at *t*=0, i.e., at the start of the simulation. The other 3 initialization conditions correspond to scenarios where there is no adaptive divergence and no barrier effect at *t*=0; however, some degree of adaptive divergence may evolve, depending on the level of initial genetic variation at adaptive loci, which is maximum for IC2, and minimum for IC4.

Figure 4 shows these results for populations of two different size, corresponding to *Ns*=2 and *Ns*=8 respectively. We see that long-term evolutionary outcomes are very similar for IC1 (high initial divergence) and IC2 (high initial standing genetic variation, but zero initial divergence). The critical migration threshold for loss of local adaptation in both cases is close to *m*_*c*,2_- the theoretically predicted critical migration threshold above which long-term divergence between habitats is not possible, regardless of initial conditions. This threshold can be read off from the solid black curves in figs. 4a and 4b— these show the theoretical predictions for the expected allele frequency, and are obtained using eqs. 8 and 4 of the main paper.

However, critical migration thresholds can be significantly lower when initial frequencies of the locally adaptive allele in the rare habitat are lower (IC3 and IC4), though in these examples, they are still higher than the LE thresholds (vertical dashed lines). Moreover, these thresholds are quite sensitive to initial levels of polymorphism in the population, which determine whether or not populations can evolve a certain minimum level of divergence rapidly enough (despite maladaptive gene flow) for LD-mediated effects to come into play, thus further reinforcing divergence. A more detailed look at simulations for IC3 and IC4 further reveals that close to the threshold for loss of local adaptation, there is considerable heterogeneity across simulation replicates— some replicates lose all polymorphism, while others evolve a high level of divergence (which is accurately predicted by the theoretical expectations, i.e., solid curve in figs. 4a and 4b). This is in contrast to the deterministic (very high *Ns*→∞) limit, where given certain initial allele frequencies, all replicates are expected to evolve towards the same equilibrium.

**Figure 4:**
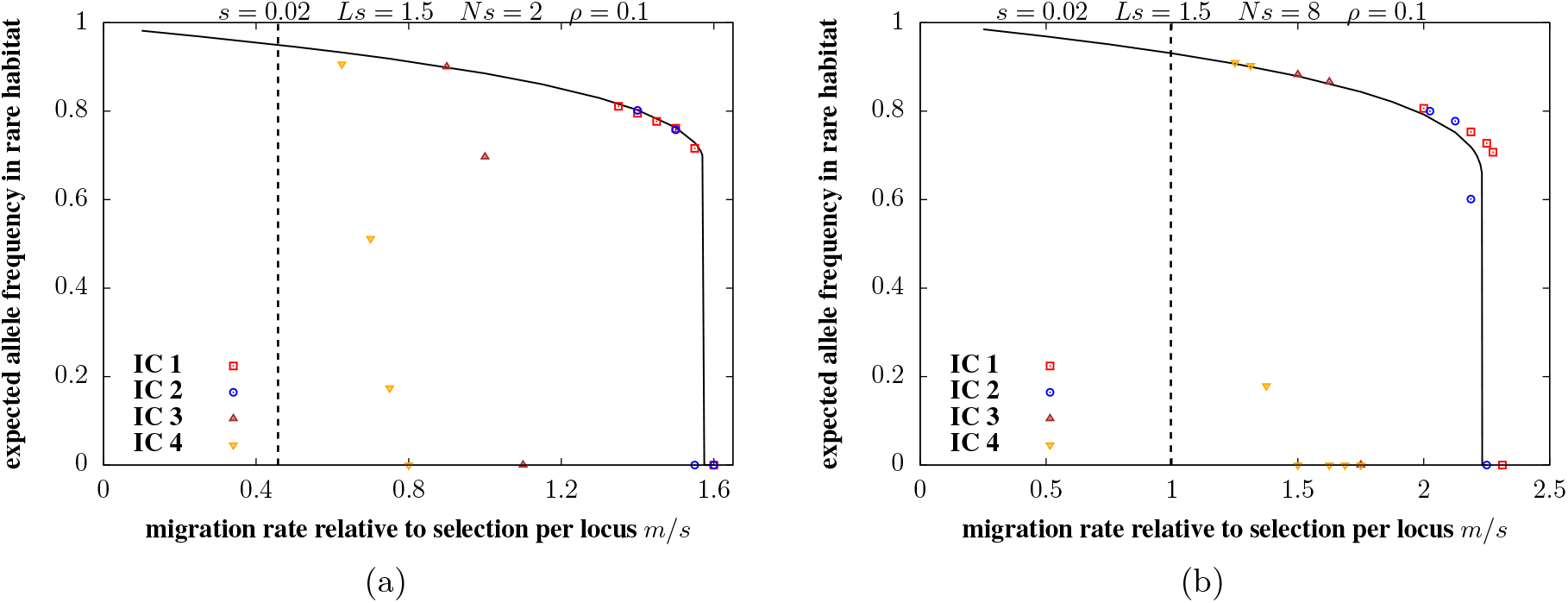
Expected frequency of locally adaptive allele in the rare habitat vs. *m/s*, the migration rate relative to selection per locus for (A) *Ns*=2 and (B) *Ns*=8. Other parameters are: *s*=0.02, *Ls*=1.5, *ρ*=0.1. The four different symbols correspond to results of individual-based simulations initialised with different allele frequencies in the rare and common habitats (see text). The solid black line shows theoretical predictions for the expected allele frequency in the rare habitat-obtained by solving for the polymorphic fixed point of eq. 8 in the main text, using eq. 4. The vertical dashed line represents the LE/single-locus critical migration threshold for a single locus (accounting for drift). Simulation results are obtained by averaging over 5 simulation replicates, each with *D*=200 demes.

